# Sigma-2 Receptor Modulators Alter Low-density Lipoprotein Receptor-mediated Lipid Uptake in Retinal Pigment Epithelial Cells

**DOI:** 10.1101/2025.08.22.670986

**Authors:** Britney N. Lizama, Aidan Reaver, Gary Look, Anthony O. Caggiano, Mary E. Hamby

## Abstract

Lipid and photoreceptor outer segment (POS) trafficking and digestion are important homeostatic functions of retinal pigment epithelial (RPE) cells, the primary cell type that degenerates in late-stage dry AMD preceded by extracellular drusen deposits of lipids and proteins. The sigma-2 receptor (S2R, TMEM97) interacts with proteins involved in lipid trafficking, such as low-density lipoprotein receptor (LDLR), which is the primary receptor for LDL-cholesterol uptake in the retina. Preclinical studies demonstrate the necessity of S2R in protecting the retina and other nervous system tissues, and loss of either S2R or LDLR has been shown to exacerbate RPE cell death and visual dysfunction. Targeting the regulatory receptors of lipid and protein trafficking functions in RPE cells represents a tractable therapeutic strategy for dry AMD. We previously demonstrated that small molecule modulators of S2R can rescue RPE POS trafficking deficits induced by exogenous stressors. Given that disruption in lipid homeostasis is a key factor in dry AMD pathogenesis and S2R interacts with LDLR, we hypothesized that LDLR-mediated lipid trafficking in RPE could be altered using S2R-targeting small molecules. In this study, RPE cells treated with S2R modulators increased LDL uptake, assessed using fluorescently-labeled LDL, compared to vehicle-treatment. Lentiviral shRNAs to reduce TMEM97 or LDLR expression, or a neutralizing blocking antibody against LDLR, reduced S2R modulator-mediated LDL uptake, confirming the requirement of TMEM97 and LDLR for S2R modulator effects. Together, these data elaborate on a potential mechanism that may underlie the favorable reduction in geographic atrophy lesion size found in participants treated with the S2R modulator zervimensine in the Ph2 clinical trial MAGNIFY in patients with geographic atrophy (COG2201, NCT05893537).

**Highlights:**

- Retinal pigment epithelial (RPE) cells express the sigma-2 receptor (S2R, TMEM97)
- Low-density lipoprotein (LDL) uptake assay in RPE cells serves as S2R functional assay
- S2R modulators, including zervimesine (CT1812), increase RPE cell LDL uptake
- S2R modulator effects are TMEM97- and LDLR- dependent

## 1. Introduction

AMD is the most common cause of blindness in people over 50 yrs of age, affecting approximately 200 million people worldwide (Fleckenstein et al., 2024). There are two forms of AMD: wet (neovascular) AMD and the more common form of the disease dry (non-neovascular) AMD. Wet AMD is characterized by maladaptive development of vasculature under the macula and is typically preceded by dry AMD. Dry AMD is characterized at the cellular level by chronic inflammation, altered lipid trafficking, accumulation of extracellular lipid-rich deposits (known as drusen), increased oxidative stress, and impaired photoreceptor function. Late-stage dry AMD, geographic atrophy, is characterized by the loss of retinal pigment epithelial (RPE) cells and progressive deterioration of the retina. To date, there are two FDA-approved treatment options for dry AMD, which provide moderate slowing of vision loss (Danzig et al., 2024; Liao et al., 2022); however the requirement for regular intravitreal injections can produce adverse effects such as retinal tearing and conversion to wet AMD (Jaffe et al., 2021; Kailani et al., 2025; Liao et al., 2022; Nadeem et al., 2023; Śpiewak et al., 2024).

The progression to GA and RPE loss is preceded by drusen formation (Veerappan et al., 2016) (Veerappan et al., 2016). Under healthy conditions, the RPE maintains efficient lipid metabolism to avoid lipid and lipoprotein accumulation. RPE cells derive cholesterol from ingestion of systemically-derived lipoproteins and from phagocytosis and recycling of photoreceptor outer segments (POS) (Pikuleva and Curcio, 2014). The RPE must maintain a balance between lipids entering the retina from exogenous sources. Large amounts of systemic lipoproteins, such as LDL, enter the RPE through the choriocapillaris endothelium, cross the Bruch’s membrane, and are internalized by the RPE (Gordiyenko et al., 2004). Cholesterol derived from LDL can be recycled back to photoreceptors or removed through normal efflux via ApoA-I and high-density lipoprotein transport. The RPE’s ability to maintain balanced lipid traffic is disrupted in dry AMD (Curcio, 2018; Curcio et al., 2011; Yamada et al., 2008), contributing to drusen formation and the aggravation of inflammation and oxidative stress. The combination of these pathophysiological manifestations leads to decline in cell health and function in the RPE and photoreceptors, ultimately causing cell death and visual impairment (Keeling et al., 2019, 2018).

Multiple genome-wide association studies (GWAS) have elucidated links between dry AMD and key cellular pathways, with single nucleotide polymorphisms (SNPs) in several loci encoding genes related to the complement pathway conferring altered risk for developing dry AMD (Arakawa et al., 2011; Fritsche et al., 2016; Helgason et al., 2013; Raychaudhuri et al., 2011; Seddon et al., 2013; Yan et al., 2018). These GWAS studies precipitated and bolstered support of drug discovery research targeting the complement pathway, which has henceforth culminated in the approvals of complement inhibitors pegcetacoplan and avacincaptad pegol, two commercially available treatments for geographic atrophy secondary to AMD (Danzig et al., 2024; Liao et al., 2022, Liao et al., 2020). In addition to complement, the GWAS studies identified AMD-risk loci that include genes related to lipid metabolism, as well as a SNP in the *TMEM97 (S2R)/VTN* locus that was associated with decreased risk for dry AMD (Fritsche et al., 2016; Yan et al., 2018), thus providing a potential genetic link of TMEM97/lipid metabolism to dAMD

In fact, several other lines of evidence – preclinical and clinical – support targeting TMEM97 and/or lipid metabolism for dAMD. Transmembrane protein 97 (TMEM97), more commonly known as the sigma-2 receptor (S2R), is a transmembrane receptor expressed in multiple cell types. S2R regulates several cell functions that are affected by degenerative diseases, including cholesterol biosynthesis/trafficking, lipid membrane-bound protein trafficking, and autophagy (Lizama et al., 2023). We have shown that small molecule modulators of the S2R, including the investigational therapeutic CT1812 (zervimesine), restore RPE POS trafficking deficits induced by amyloid-beta oligomer (Aβ42) and hydrogen peroxide *in vitro* (Lizama et al., 2025). Zervimesine treatment has conferred benefits in clinical trials and in models of CNS degeneration (Izzo et al., 2021; van Dyck et al., 2024; Vijverberg et al., 2024, 2025). Further, zervimesine has been tested in a phase 2 clinical trial for geographic atrophy (GA) secondary to dry AMD (MAGNIFY, COG2201, NCT05893537), with encouraging results showing GA lesion size in zervimesine-treated participants were 28.2% smaller compared to placebo at 18 months (Cognition Therapeutics Inc., 2025).

Evidence that S2R is mechanistically immersed in pathways impacted by degenerative disease is also found in preclinical lipid trafficking studies. Pairwise proximity ligation assays in Hela cell knockout models (Riad et al., 2018) and human brains (Riad et al., 2020) have confirmed the interaction between the LDL receptor (LDLR), the sigma-2 receptor (TMEM97, S2R) and its coreceptor PGRMC1. The S2R-LDLR complex is involved in Aβ42 uptake, as supported by *in vitro* evidence in which TMEM97 knockout reduced Aβ42 uptake in human cancer HeLa cells (Riad et al., 2020). Additionally, pharmacological modulators of S2R dampened uptake of Aβ42 in primary rat neurons (Riad et al., 2020) and restored vesicular trafficking deficits incurred by Aβ42 oligomers (Izzo et al., 2014a, 2014b). Rapid internalization of LDL cholesterol by LDLR has also been shown to require S2R, as genetic knockout of either TMEM97 or PGRMC1 in HeLa cell lines decreased the rate of internalization of ^3^H-labeled LDL and fluorescently tagged LDL (Riad et al., 2018). These data show potential for S2R modulation as an effective method for affecting this trafficking. Moreover, this growing body of research begs the question whether S2R could be a promising therapeutic target for AMD due to its interactions with the LDLR and important role in lipid trafficking. It has been shown that LDLR is the primary receptor for RPE LDL-cholesterol uptake (Go and Mani, 2012; Tserentsoodol et al., 2006), and that loss of LDLR exacerbates and contributes to dry AMD-like phenotypes (Chen et al., 2009; Sreekumar et al., 2022). Multiple GWAS studies have linked the lipid metabolism pathway to AMD (Fritsche et al., 2014, 2013; Neale et al., 2010), and there is an age-dependent increase in subretinal lipid/protein deposits (Huisingh et al., 2016) that, in in vivo models of exogenous or genetic AMD risk factors – such as cigarette smoke exposure (Espinosa-Heidmann et al., 2006) or mutations in AD-linked genes that increase retinal Aβ burden (Lynn et al., 2025) – are thought to contribute to POS trafficking dysfunction and retinal degeneration. LDLR^-/-^ mice exhibited impaired retinal function and lipid accumulation in the RPE, and RPE and photoreceptor degeneration was exacerbated when oxidative stress was induced via NaIO_3_ (Sreekumar et al., 2022). Degeneration of Bruch’s membrane accompanied by accumulation of lipid particles was also reported after examining retinas taken from LDLR^-/-^ mice by transmission electron microscopy (TEM) (Rudolf et al., 2004).

Given the lipid dysfunction critically linked with cell demise in AMD, and the known links between the disease, TMEM97, and LDLR, we hypothesized that S2R-targeting small molecules would affect lipid uptake in RPE. To this end, we developed an *in vitro* assay to assess the ability of proprietary small molecule modulators of the S2R to alter lipid uptake, leveraging learnings from prior functional and structural interaction of TMEM97 and LDLR studies that have been performed in heterologous cell types (Riad et al., 2020, 2018). Using differentiated ARPE-19 cultures, we confirm the presence of the S2R components and LDLR in differentiated RPE cells and the ability of these cells to robustly uptake LDL via imaging of fluorescently tagged LDL. With this optimized assay, we demonstrate for the first time the ability of S2R modulators to increase LDL uptake in RPE cells, and provide mechanistic support of a dependency of LDLR in the ability of zervimesine and other S2R modulators in mediating this effect.

## 2 Methods

### 2.1 Cell Culture

ARPE19 cells were purchased from ATCC (70079881), expanded, and frozen at passage 5. Experimental cells were all thawed and cultured from passage 6 in T75 flasks in DMEM-F12 supplemented with 10% FBS and 1% Pen/Strep. For experiments, cells were seeded into laminin-coated 96-well plates at 25,000 cells/well and grown in “MEM-Nic” - MEM supplemented with 1% FBS, 1% Pen/Strep, 1% GlutaMax, 1% N2, 20 ng/mL hydrocortisone, 0.013 ng/mL triiodo-L-thyronine, 0.25 mg/mL taurine, and 10 mM nicotinamide (Hazim et al., 2019). Cells were grown in MEM-Nic for two weeks prior to experiments to induce differentiation and polarization, and experiments were performed between passage 8-12. For studies assessing differentiation, cells were imaged and RNA was harvested after one or two weeks of culturing with either DMEM-F12 or MEM-Nic feeding medium (Supp Fig 1). The progression of differentiation was confirmed by qRT-PCR of RPE-specific markers *BEST1, MERTK*, and *RPE65* (Hazim et al., 2019). For TMEM97 or LDLR knockdown studies, lentiviral vectors were introduced after 7-10 days of differentiation, and cells were kept in culture with regular feeding every 2-3 days with MEM-Nic for seven days thereafter. For experiments assessing LDL uptake, S2R modulator or U18 treatments were initiated between 14-17 days of differentiation, with treatment lasting 16 hr before initiation of LDL uptake. LDL uptake occurred with S2R modulator/U18 concentrations maintained in the media, was started directly after 16 hr pretreatment, and endpoint assays were performed 4 hr after LDL was added (LDL uptake assay, mRNA and protein harvest, cell fixation).

**Figure 1.**
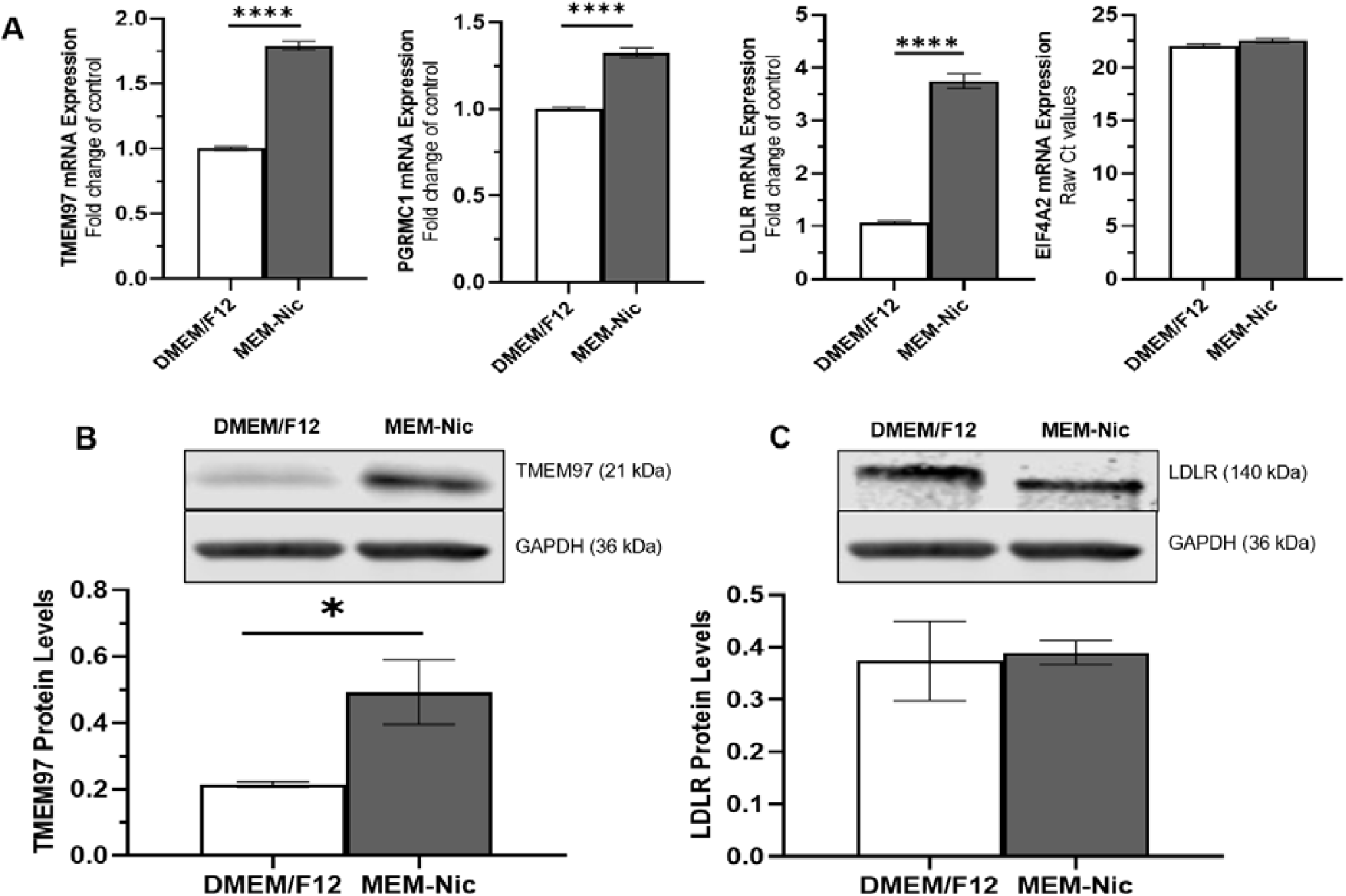
S2R components are expressed in mature RPE cells. **A)** qRT-PCR analysis of TMEM97 and co-receptor expression after 14 days of differentiation (MEM-Nic) compared to control (DMEM/F12), control conditions n=14, DIV14 conditions n=10, normalized to EIF4A2 + control, mean ± SEM, unpaired t-test, **** p≤0.0001. Representative western blots of TMEM97 (**B)**, LDLR **(C)**, and housekeeping protein GAPDH in ARPE19 cells after 14 days of differentiation (MEM-Nic) compared to control (DMEM/F12). Densitometry of TMEM97 **(B)** and LDLR **(C)** normalized to GAPDH plotted, mean ± SEM of N=4; unpaired t-test,* p≤ 0.05.

Wildtype and TMEM97 KO HEK-293T cells were purchased from Abcam (ab266499) and sub-cultured in T75 flasks in DMEM supplemented with 10% FBS and 1% Pen/Strep. For experiments, cells were seeded into laminin-coated 96-well plates at 15,000 cells/well and incubated for 24 hr prior to treatments. Experiments were performed using cultures with passage <10.

### 2.2 S1R and S2R Modulators

Cognition Therapeutics S2R modulators from three chemically distinct series, CT2074, CT2168, and CT1812 (also known as the investigational therapeutic, zervimesine), were stored at room temperature as lyophilized powders until experimentation. CT1812, CT2074, and CT2168 exhibit similar S2R binding affinity in a S2R binding assay (Donkor et al., 2024; Lizama et al., 2025) and demonstrate similar pharmacology in *in vitro* functional assays using primary neurons (Izzo et al., 2014a, 2014b; Lizama et al., 2023) and RPE cells (Lizama et al., 2025). Affinities of these modulators at the S2R are in the low nanomolar range: CT1812 (Ki = 8.5 nM), CT2074 (Ki = 21 nM), and CT2168 (Ki = 1.4 nM). For CT1812, selectivity was >⍰100-fold over S2R on a radioligand binding inhibition panel from Eurofins (France), screened against 118 proteins. For CT2168, selectivity was >⍰100-fold over S2R on a radioligand binding inhibition panel from Eurofins, screened against 116 proteins. For CT2074, selectivity was >⍰100-fold over S2R on a radioligand binding inhibition panel from Eurofins, screened against 117 proteins, except for two with >⍰60-fold. S2R modulators were reconstituted in anhydrous DMSO (Merck, UK) and then diluted with culture medium to final assay concentrations.

Similarly, sigma-1 receptor (S1R) literature standards NE-100 (Tocris, #3133; S1R Ki = 1 nM, >190 fold over S1R), and PRE-084 (Tocris, #0589; S1R Ki = 52.4 nM, >190 fold over S1R) were tested at EuroFins versus the human sigma receptors, and stored at room temperature until further experimentation.

### 2.3 LDL-DyLight 550 Assay

LDL DyLight 550 (Cayman, 10011229) was diluted in serum-free media (MEM-Nic without FBS for ARPE-19; DMEM without FBS +0.3% BSA for HEK-293T) at a working concentration of 20 µg/mL and filtered with a 0.45 µm PES filter. S2R modulators were dissolved in DMSO and diluted in media (DMSO final concentration in assay wells was 0.1%). Differentiated ARPE19s were incubated with S2R modulators for 16 hr at 37°C, after which medium was removed and replaced with LDL-DyLight550 (hereafter, referred to as LDL-DyLight) working solution. S2R modulators were re-introduced with LDL-DyLight medium exchange, so that concentrations of S2R modulators remained constant. After incubation with LDL for 4 hr, cells were washed twice with 1x DPBS and fixed with 4% PFA in DPBS for 10 min at RT. Cells were washed twice after fixing with PBS and stained with 10 µg/mL Hoechst 33342 in PBS for 10 min at RT. Hoechst was removed from the cells with 2x 5 min washes with DPBS. LDL DyLight550 (540/570) and Hoechst (350/461) staining were imaged at 20x magnification using a CellInsight CX7 HCS Platform. LDL-DyLight fluorescence was quantified as fold change, normalized to DMSO vehicle control.

### 2.4 LDLR Blocking Antibody

Differentiated ARPE19s were incubated with S2R modulators for 16 hr at 37°C, after which medium was removed and replaced with half the original well volume of LDLR blocking antibody (AF2148; 1-50 µg/mL) diluted in serum-free media (MEM-Nic without FBS) for 30 min at 37°C. S2R modulators were re-introduced with the medium exchange, so that concentrations of S2R modulators remained constant. After blocking, LDL-DyLight working solution was introduced with S2R modulators and blocking antibody to maintain assay concentrations. LDL-DyLight was diluted to a 2x working concentration of 40 µg/mL and a half-well volume of LDL-DyLight working solution was added on top of the blocking media. After 4 hr incubation with LDL at 37°C, cells were washed twice with 1x DPBS and fixed with 4% PFA in DPBS for 10 min at RT. Cells were washed twice with PBS and stained with 10 µg/mL Hoechst 33342 in PBS for 10 min at RT. Hoechst was removed from the cells with 2x 5 min washes with DPBS. LDL DyLight550 (540/570) and Hoechst (350/461) staining were imaged at 20x magnification using a CellInsight CX7 HCS Platform. LDL-DyLight fluorescence was quantified as fold change, normalized to DMSO vehicle control.

### 2.5 Immunocytochemistry

Cell cultures were fixed in 4% paraformaldehyde for 10 min at room temperature and washed in 1x DPBS. For assessment of RPE marker ZO-1, cells were permeabilized with 0.1% triton-X 100, washed, and blocked with 1% BSA +0.1% Tween for 1 hr. Primary antibody was added at a 1:100 dilution in blocking buffer solution (1% BSA, 0.1% Tween) overnight, followed by washes and incubation with Alexa Fluor 647 secondary antibody (1:500) for 1 hr. After additional washes, cell nuclei were stained with Hoechst.

For assessment of LDLR, cells were not permeabilized to preserve surface plasma membrane levels of the LDLR. Immunocytochemistry was performed as above with without use of detergents in all incubation and washing steps. Cultures treated with LDLR blocking antibody (described above) were assessed for LDLR levels. After LDL-DyLight fluorescence was captured, cells treated with LDLR blocking antibody were blocked in 1% BSA for 1 hr and incubated with Alexa Fluor 647 secondary antibody (1:500, Jackson Labs) to visualize external LDLR. LDL DyLight550 (ex/em: 540/570), Hoechst (350/461), and external LDLR (651/667) staining was captured at 20x magnification using a CellInsight CX7 HCS Platform.

### 2.6 Fluorescence Imaging Analysis

LDL-DyLight 550 and Alexa Fluor 647 immunofluorescence were quantified using the CellInsight CX7 HCS Platform’s Spot Detection software. Hoechst 33342 staining was used to identify individual cells, with parameters set to exclude cell clusters in which nuclei could not be segmented, or Hoechst-positive staining <20 µm^2^. A mask was projected to the edges of the cytosol surrounding each nucleus. Fluorophore spot intensity per cell was measured, with parameters for spot size and intensity set to avoid detection errors caused by background or aberrant fluorescence. For each experimental plate, spot intensity per cell was recorded as an average of 16 fields per well.

### 2.7 RNA Isolation and RT-qPCR

Following treatment, media was removed from the wells and cells were washed twice with PBS. Cells were lysed with 150 µL Buffer RLT (Qiagen, 74106) for 3 min at RT. Lysate was transferred to tubes and stored at -80°C. RNA concentrations were measured using a NanoDrop One/One Microvolume UV-Vis Spectrophotometer. 0.6 µg RNA was used to generate cDNA with the SuperScript IV First-Strand Synthesis System (Fisher, 18-091-200) according to the manufacturer’s protocol. RT-qPCR was performed using Taqman Fast Advanced Master Mix and Taqman gene expression assays (Thermo, 4331182: Hs00756996_g1, Hs00299877_m1, Hs01092524_m1, Hs00427620_m1, Hs00264835_m1. Applied Biosystems, 4325792). RT-qPCR plates were quantified using a QuantStudio 5 Real-Time PCR System. Ct values measured were normalized to housekeeping gene (GAPDH or EIF4A2, indicated within each figure) and experimental vehicle controls (ΔΔCT), and converted to fold change (2^-ΔΔCT).

### 2.8 Protein Isolation

Following treatment, media was removed from the wells and cells were washed with PBS. Cells were lysed with M-PER Buffer (Thermo, 78501) spiked with a protease inhibitor cocktail (Thermo, 78440) for 5 min at RT. Wells were scraped and lysate was transferred to tubes to incubate on ice for 10 min. Lysates were centrifuged at 10,000 x g for 10 min. Lysate supernatant and pellet were separated, flash frozen using liquid nitrogen, and stored at -80°C for western blot analysis.

### 2.9 Western Blot

Protein lysate (20 µg) was diluted 4:1 with 4x Laemmli sample buffer (Bio-Rad, 161-0747) and heated to 95°C for 10 min. Samples were loaded onto a 4-15% Tris-HCL gel (Bio-Rad, 345-0028) and ran at 125 V for 75 min. Protein was transferred onto a nitrocellulose membrane (Bio-Rad, 162-0112) and blocked using Intercept Blocking Buffer for 1 hr. The membrane was incubated in primary antibody overnight (Cell Signaling, 62790, 1:100. Abcam, ab8245, 1:2000. Cell Signaling, 139015, 1:5000. Abcam, ab52818-1001, 1:1000). The membrane was then washed 3 times with 1x TBST for 5 min and stained with secondary antibody (Licor, 926-32210 and 925-32211, 1:10,000) for 1 hr, protected from light. After washing the membrane three times, 5 min each with 1x TBST, protein bands were imaged using a LICOR Odyssey (ex/em: 778/795). Densitometry was performed using Image Studio. Protein levels were normalized to GAPDH or Vinculin, and DMSO vehicle controls.

### 2.10 Lentiviral transduction

The lentiviral vectors targeting TMEM97 (3LVM(LVshRNA)-C-27346-5) and LDLR (3LVM(LVshRNA)-C-3949-5) were purchased from Vector Builder. ARPE-19 cells were differentiated in 96-well plates in MEM-Nic medium. Within each plate, cultures were assigned into four groups: untreated control, non-transduced control treated with polybrene (10 μg/mL final) only, transduced with a negative control lentiviral vector (scrambled shRNA), and transduced with a 1:1:1 mixture of three shRNA vectors targeting TMEM97 (shTMEM97) or LDLR (shLDLR). Cells were transduced overnight with scrambled control or shTMEM97 lentiviral vectors at a multiplicity of infection (MOI) of 9, or shLDLR vectors at MOI of 3, after which culture medium was fully changed. Cells were further incubated for up to 7 days after transduction, with regular medium changes every 2-3 days, and then treated with S2R modulators for 16 hr. Cells in sister plates within each experiment were either harvested for protein or mRNA, fixed for immunocytochemistry, or used for the LDL uptake assay. TMEM97 or LDLR knockdown was confirmed by qRT-PCR and by western blotting. Protein, mRNA, and LDL-DyLight levels from each treatment group were normalized to polybrene-treated cultures within each experiment.

### 2.11 Statistical analyses

Statistical method and normalization details are indicated within each figure legend. Analysis of the data was performed using either an unpaired t test, one-way analysis of variance (ANOVA), or two-way ANOVA using PRISM software (version 10.3.1; GraphPad Software Inc, Boston, MA, USA). One-way ANOVA analyses were accompanied with Dunnett’s multiple comparisons test, and two-way ANOVA analyses were accompanied with Tukey’s multiple comparisons test. Data are expressed as mean ± SEM. In all analyses, p ≤ 0.05 was considered statistically significant.

## 3 Results

### 3.1 Characterization of Differentiated ARPE-19 cells: Differentiation markers and S2R expression

Several reports indicate that appropriate differentiation of ARPE-19 cells allows for a more physiological phenotype of cells resembling native RPE cells in their typical capacity to phagocytose and degrade photoreceptor outer segments. Consistent with prior knowledge, undifferentiated ARPE-19 cultures exhibit fibroblast-like morphology, even when cultured for 14 days *in vitro* (DIV14) (Supplementary Figure 1A). Unlike undifferentiated cultures, ARPE-19 cells differentiated from DIV1 to DIV14 form a compact monolayer, exhibit canonical cobblestone morphology (Supplementary Figure 1A) and are zonula occludens-1 (ZO-1, tight junction protein) positive (Supplementary Figure 1B). Compared to undifferentiated control cells, differentiated ARPE-19 cells express higher transcript levels of mature RPE-specific markers bestrophin-1 (BEST1, 1384-fold increase ± 56.23), c-Mer tyrosine kinase (MERTK, 6-fold increase ± 0.14), and retinoid isomerohydrolase RPE65 (RPE65, 765-fold increase ± 35.89) (Supplementary Figure 1C). Transcript levels of S2R complex components transmembrane protein 97 (TMEM97) and progesterone receptor membrane component 1 (PGRMC1) were significantly higher in differentiated versus undifferentiated control RPE cells (TMEM97: 1.8-fold increase ± 0.034; PGRMC1: 1.3-fold increase ± 0.028) (Figure 1A). TMEM97 protein levels reflect the increase in mRNA expression levels, with a 2.3-fold increase ± 0.097 in TMEM97 protein levels in differentiated versus control RPE cells (Figure 1B). LDLR mRNA levels were also significantly higher in differentiated versus control RPE cells (3.7-fold increase ± 0.14) (Figure 1A); however, there was no significant change in total LDLR protein levels observed in differentiated versus control RPE cells (Figure 1C).

### 3.2 S2R modulators increase LDL uptake in RPE cells

Given that S2R and LDLR are expressed in differentiated ARPE-19 cells and their interaction has been shown to assist in LDL uptake in cancer cells (HeLa), a fluorescence-based LDL (LDL tagged with a DyLight fluorophore (i.e., LDL-DyLight)) uptake assay was developed in differentiated ARPE-19 cells. To optimize the assay, a commercially available small molecule, U18666A (hereafter referred to as U18), which is known to robustly trigger LDL uptake via Niemann-Pick’s protein C 1 (NPC1), was used as a positive control. Treatment with U18 for 16 hr prior to and during LDL-Dylight addition increased levels of LDL-DyLight in a concentration-dependent manner when assessed 4 hr post-LDL-Dylight addition: 1 μM U18 induced a 1.9-fold (± 0.10) increase, 5 μM induced a 3-fold (± 0.22) increase, 10 μM induced a 4.1-fold (± 0.28) increase, and 30 μM induced a 5.1-fold (± 0.30) increase over vehicle control (EC_50_ = 4.53 µM) (Figure 2A*i*). Images procured via fluorescence microscopy demonstrate LDL-DyLight puncta in vehicle-treated control cells in 100% of cells at 30 µM (Figure 2A*ii*, left panels) that is highly increased in cultures treated with U18 (30 µM) for 16 h prior to LDL-Dylight addition (Figure 2A*ii*, right panels). We next assessed whether the U18-mediated increase in LDL was related to a corresponding increase in total LDLR levels induced by U18. While LDLR transcript levels increased after 16 hr U18 treatment in a concentration-dependent manner (Figure 2B*i*), and LDLR mRNA was significantly positively correlated with LDL-DyLight levels (r = 0.86; Figure 2B*ii*), no increase in total LDLR protein levels, as assessed via western blot, was observed with any concentration of U18 (Figure 2B*iii*), and no correlation of LDL-DyLight levels to LDLR total protein levels was observed (Figure 2Biv). In contrast, both TMEM97 mRNA transcript and protein levels increased with U18 treatment in a concentration-dependent manner, with levels peaking at 10-30 µM (Fig C*i, iii*). However, similar to LDLR, a significant positive correlation was observed between LDL-DyLight levels and TMEM97 mRNA levels (r = 0.89) but not TMEM97 protein levels (Figure 2C*ii, iv*). Three chemically distinct S2R modulators were tested for their ability to modulate LDL uptake in differentiated RPE cells using the LDL-DyLight assay (Figure 3A*i, ii*) and showed comparable increases in LDL levels after 4 hr with compound pretreatment (16 hr; 10 µM). In contrast to S2R-targeting small molecules, commercially available sigma-1 receptor (S1R) tool compounds NE-100 and PRE-084 did not significantly alter LDL levels (Figure 3B).

**Figure 2.**
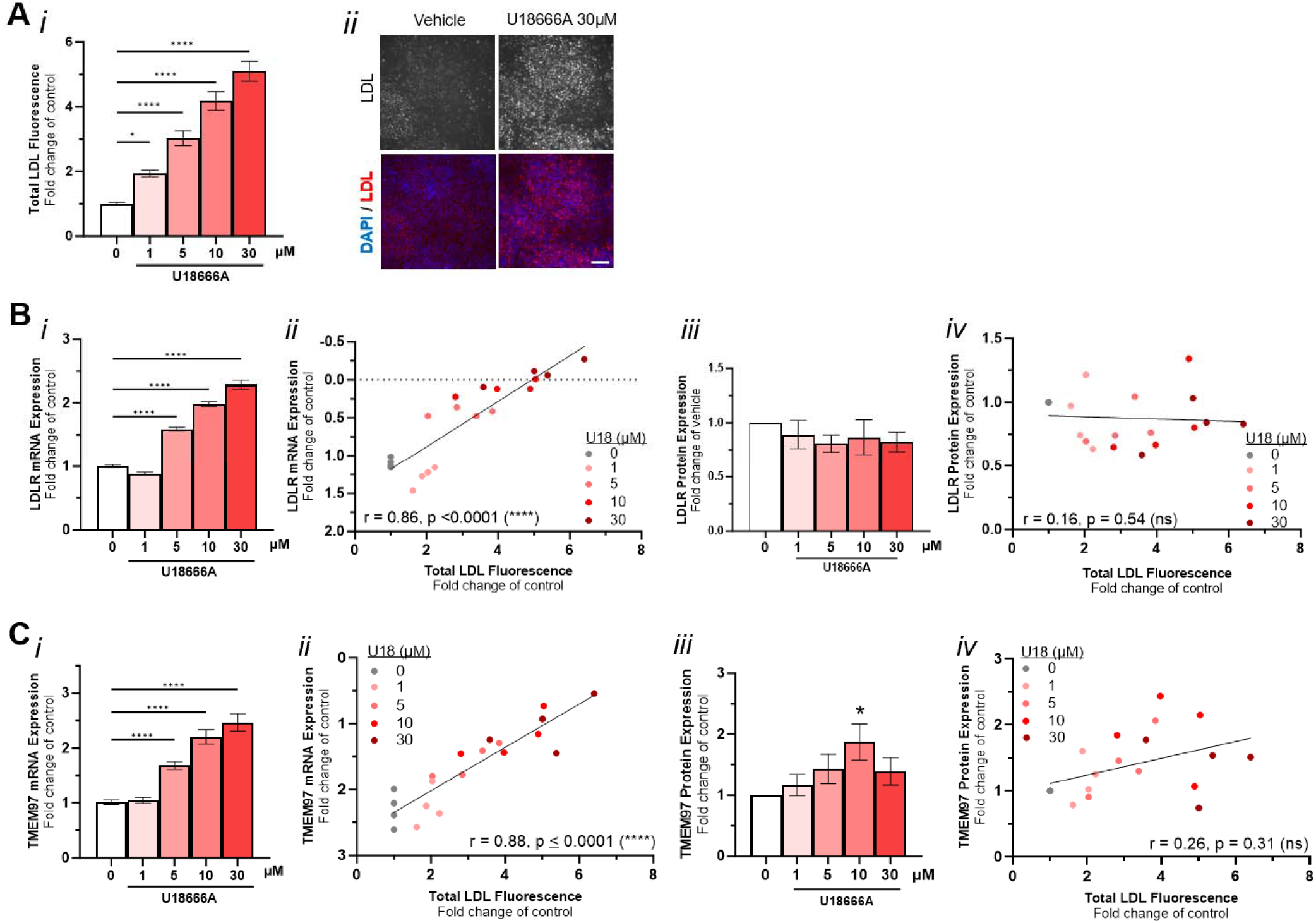
LDL uptake assay optimized in mature RPE cells using well-characterized pharmacological tool U18666A. **A) i**. Quantification of LDL-DyLight total intensity per cell (assessed using CX7 Spot Detector) after treatment with increasing concentrations of U18, normalized to vehicle, mean ± SEM of N=4; one-way ANOVA, Dunnett’s multiple comparison test; **** p≤0.0001, U18 vs control (0). *ii*. LDL-DyLight assay fluorescence representative images of differentiated ARPE-19 cells treated with U18666A. 20x magnification, NA 0.70; scale bar = 50µm. **B)** *i*. qRT-PCR analysis of LDLR after treatment with ascending concentrations of U18, normalized to EIF4A2 and vehicle, mean ± SEM of N=4, one-way ANOVA, Dunnett’s multiple comparison test. *ii*. Corresponding correlation analysis comparing the expression of mRNA (ΔCT, normalized to EIF4A2) with LDL-DyLight fold change, **** p≤0.0001. *iii*. Densitometry of western blots for LDLR after treatment with ascending concentrations of U18, normalized to GAPDH and vehicle, mean ± SEM of N=4, one-way ANOVA, Dunnett’s multiple comparison test.. *iv*. Corresponding correlation analysis of fold change in protein levels with LDL-DyLight fold change. **C)** *i*. qRT-PCR analysis of TMEM97 after treatment with ascending concentrations of U18, normalized to EIF4A2 and vehicle, mean ± SEM of N=4, one-way ANOVA, Dunnett’s multiple comparison test. *ii*. Corresponding correlation analysis comparing the expression of mRNA (ΔCT, normalized to EIF4A2) with LDL-DyLight fold change, **** p≤0.0001. *iii*. Densitometry of western blots for TMEM97 after treatment with ascending concentrations of U18, normalized to GAPDH and vehicle, mean ± SEM of N=4, one-way ANOVA, Dunnett’s multiple comparison test. *iv*. Corresponding correlation analysis of fold change in protein levels with LDL-DyLight fold change, * p≤0.05.

**Figure 3.**
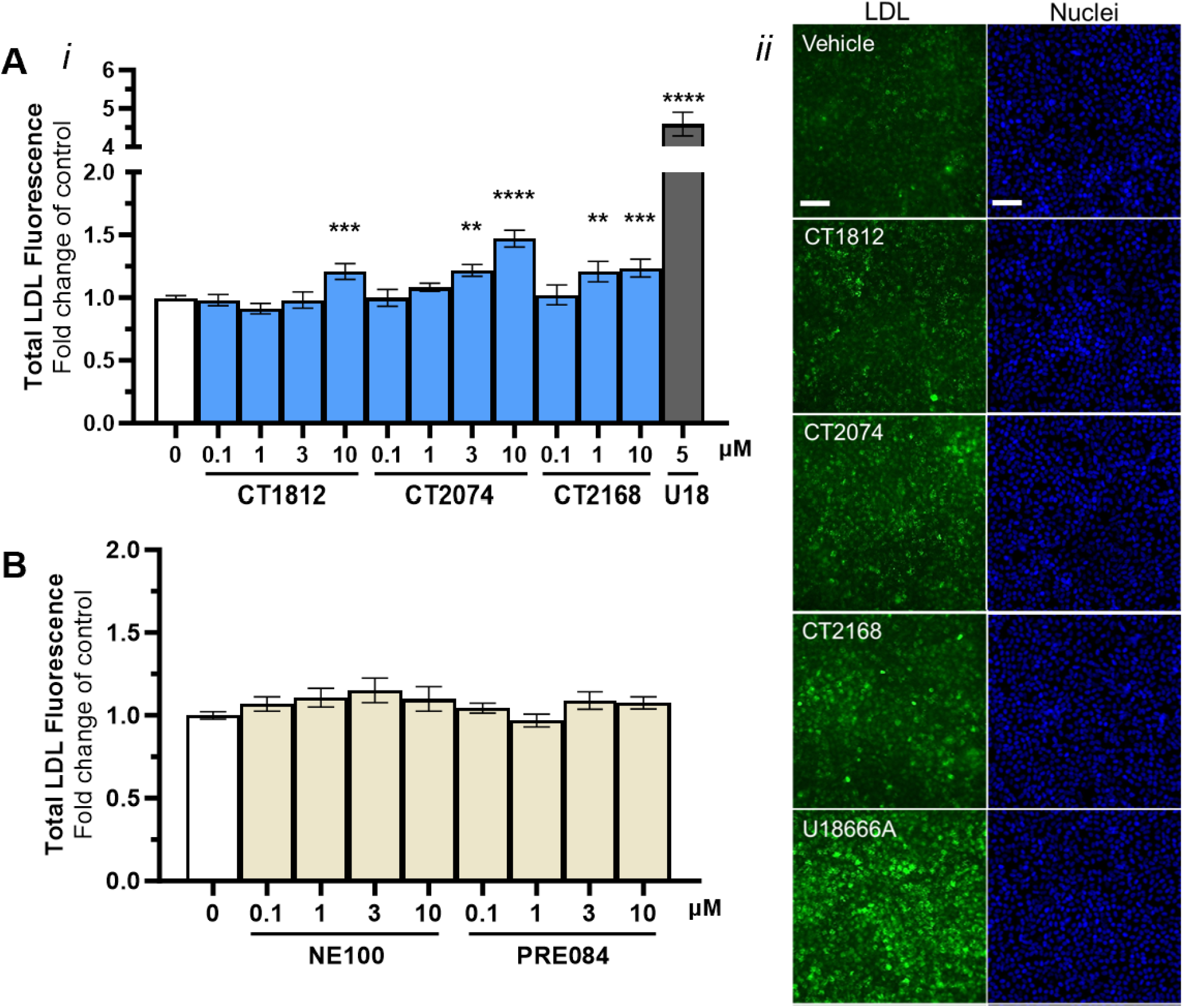
LDL uptake in RPE cells is S2R modulator-specific. **A)** *i*. Quantification of LDL-DyLight total intensity per cell (assessed using CX7 Spot Detector) in differentiated ARPE-19 cells treated with S2R modulators (CT1812, CT2074, CT2168), normalized to vehicle (0), mean ± SEM of N=5, one-way ANOVA, Dunnett’s multiple comparison test; ** p≤0.0021, *** p≤0.0002, **** p≤0.0001 treatment vs vehicle. *ii*. Representative images of LDL-Dylight fluorescence (green) and nuclei stain Hoechst (blue) after treatment with vehicle, U18 (5µM) or S2R modulators (CT1812, CT2074, CT2168 at 10µM). 20x magnification, NA 0.70; Scale bar = 50µm. **B)** Quantification of LDL-DyLight total intensity per cell as assessed in A after treatment with S1R-selective modulators NE-100 and PRE-084, normalized to vehicle control (0), mean ± SEM of N=3, one-way ANOVA, Dunnett’s multiple comparison test.

### 3.2 Increase in LDL uptake by S2R modulators requires LDLR expression

Treatment with S2R modulators CT1812 (zervimesine), CT2074 and CT2168 (16 hr) did not significantly change total LDLR mRNA nor total protein levels (Figure 4A*i, ii*), thus verifying a change in total expression levels did not underlie the S2R-mediated increase in LDL uptake. Given the pharmacological effect on LDL uptake was similar across these S2R modulators tested, (Figure 3) and other previously explored functions in RPE cells (Lizama et al., 2025), CT2074 was selected to further investigate the S2R modulator-mediated increase in LDL uptake mechanism of action. Next, whether LDL uptake was LDLR-dependent, under basal and S2R modulator-treated conditions, was investigated. Under basal conditions, an LDLR-selective blocking antibody was incubated with vehicle-treated cells prior to (30 min) and during the LDL uptake assay (4 hr) (Figure 4B). A significant reduction in LDL-DyLight fluorescence (30 µg/mL of LDLR blocking antibody: 0.37-fold of vehicle control ± 0.18; IC_50_ = 9.8 µg/mL) (Figure 4B*i*) was observed when LDLR was blocked (Figure 4B*ii*).

**Figure 4.**
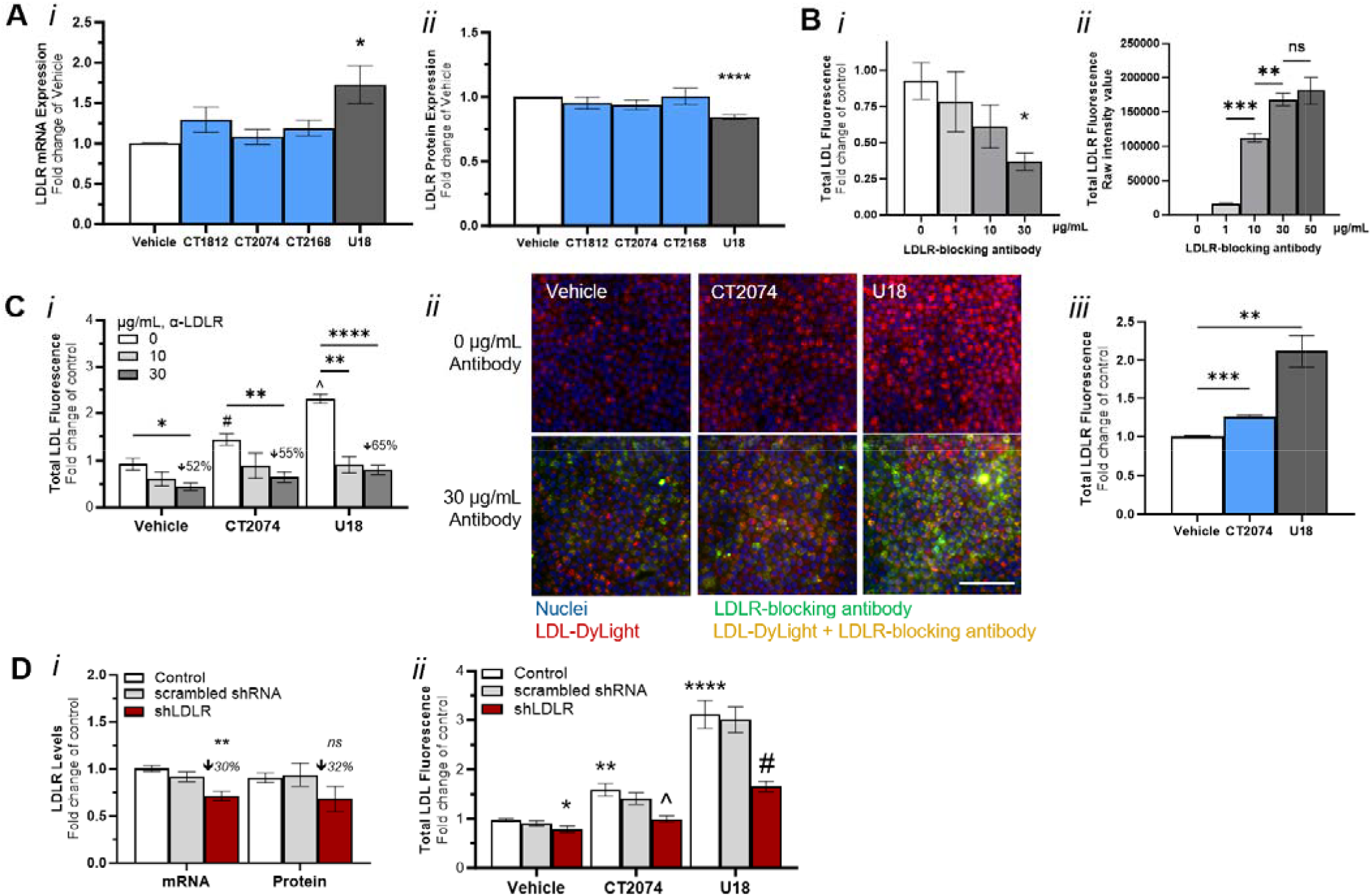
S2R modulator increases in LDL uptake require LDLR. **A)** qRT-PCR analysis (*i*.) of LDLR mRNA normalized to GAPDH and vehicle, and western blot densitometry (*ii*.) of LDLR protein normalized to Vinculin and vehicle, after treatment with S2R modulators (10 µM) or U18 (5 µM). Mean ± SEM of N=5, one-way ANOVA, Dunnett’s multiple comparison test; * p≤0.05, **** p≤0.0001. **B)** *i*. Quantification of LDL-DyLight total intensity per cell (assessed using CX7 Spot Detector) in differentiated ARPE-19 cells treated with LDLR blocking antibody, normalized to vehicle, mean ± SEM of N=3, one-way ANOVA, Dunnett’s multiple comparison test; * p≤0.05. Calculated IC_50_ = 9.8 µg/mL. *ii*. LDLR-positivity was assessed using Alexa647-conjugated secondary antibody targeting an LDLR blocking antibody (raw fluorescence intensity per cell assessed using CX7 Spot Detector), mean ± SEM of N=3, one-way ANOVA, Dunnett’s multiple comparison test; **p≤0.01, ***p≤0.001.**C)** *i*. Quantification of LDL-DyLight fluorescence after treatment with vehicle, CT2074 (10µM) or U18 (5µM) and 0, 10, or 30 µg/mL LDLR blocking antibody, normalized to vehicle, mean ± SEM of N=3, two-way ANOVA, Dunnett’s multiple comparison test; * p≤0.05, ** p≤0.01, **** p≤0.0001, 0 µg/mL vs 30 µg/mL within treatment condition; # p≤0.05 CT2074 vs vehicle + 0 µg/mL, ^ p≤0.0001 U18 vs vehicle + 0 µg/mL. *ii*. Representative images of ICC performed on non-permeabilized cells: LDL-DyLight (red), LDLR blocking antibody (30µg/mL, green), and Hoechst-positive nuclei (blue). 20x magnification, NA 0.70, scale bar = 50µm. iii. LDLR-positivity was assessed as in B, after treatment with 30 µg/mL LDLR blocking antibody, mean ± SEM of N=3, one-way ANOVA, Dunnett’s multiple comparison test; **p≤0.01, ****p≤0.0001. **D)** *i*. LDLR levels in differentiated ARPE-19 cells treated with lentiviral vector carrying shRNA targeting LDLR (shLDLR) or control vector (scRNA) were assessed by qRT-PCR (left) analysis of LDLR mRNA (normalized to EIF4A2 and polybrene treatment) and by immunofluorescence staining of LDLR protein (right) (fluorescence intensity per cell normalized to polybrene treatment), mean ± SEM of N=3, with percent LDLR knockdown indicated. *ii*. Quantification of LDL-DyLight fluorescence after treatment with vehicle, CT2074 (10µM) or U18 (5µM) in control cells (white) or after treatment with scRNA (gray) or shLDLR (red) lentiviral vectors, normalized to vehicle, mean ± SEM of N=4, two-way ANOVA, Dunnett’s multiple comparison test; **p≤0.01 Control CT2074 vs vehicle, ****p≤0.0001 Control U18 vs vehicle, ^p≤0.05 CT2074 shLDLR vs control, #p≤0.01 U18 shLDLR vs control.

To test the hypothesis that S2R modulator-mediated LDL uptake was LDLR-dependent, LDL uptake was assessed in the presence or absence of the LDLR blocking antibody (with 30 min pre-treatment). In the absence of the blocking antibody, CT2074 significantly increased LDL uptake compared to control (1.44-fold change of vehicle ± 0.13); however, in the presence of the LDLR blocking antibody (CT2074 + 30 µg/mL antibody) a significant 55% reduction in uptake was seen (0.65-fold of vehicle control ± 0.11) (Figure 4C*i*). Similarly, treatment with the LDLR-specific blocking antibody induced a significant 65% decrease in U18-mediated LDL-DyLight fluorescence (U18 + 0 µg/mL antibody: 2.30-fold of vehicle control ± 0.10; U18 + 30 µg/mL antibody: 0.81-fold of vehicle control ± 0.10) (Figure 4C*i*). Whether LDLR surface levels were affected with S2R modulators was next assessed via immunofluorescence detection of the LDLR blocking antibody in non-permeabilized fixed cells. Fluorescence microscopy images of non-permeabilized cells show basally (vehicle-treated) low levels of surface, plasma membrane-localized LDLR (Figure 4C*ii*, green; left panel), levels which were increased in CT2074-(middle panel, green) and U18-treated cells (right panel, green), corresponding to an increase in LDL-DyLight fluorescence (red) in CT2074-(middle panel, red) and U18-treated cells (right panel, red) (Figure 4C*ii*). This effect was quantified and CT2074-treated cells exhibited significantly higher levels of LDLR-positive fluorescence compared to vehicle (1.26-fold change of vehicle ± 0.02) (Figure 4C*iii*, blue bar). U18-treated cells also showed high levels of LDLR-positive fluorescence (2.2-fold change of vehicle ± 0.16) (Figure 4C*iii*, gray bar).

To further confirm whether LDLR is necessary for S2R-mediated LDL uptake, lentiviral vectors targeting LDLR (shLDLR) were used. In shLDLR transduced cultures, LDLR mRNA was significantly decreased by 30% (0.71-fold of control ± 0.05) compared to control, while scrambled control shRNA (scRNA) did not significantly change LDLR mRNA levels (Figure 4D*i*, left). To determine whether LDLR protein levels, specifically at the plasma membrane surface, were successfully decreased, immunocytochemistry was conducted on fixed, non-permeabilized cells using the LDLR-specific antibody used in Figure 4B-C. Similar to the observed decrease in LDLR mRNA, LDLR protein levels in shLDLR-treated cultures were decreased by 32% (0.68-fold of control ± 0.13) compared to control (Figure 4D*i*, right), although the difference was not statistically different (p=0.33). We next tested whether shLDLR-treated cultures would also exhibit a decrease in LDL uptake. At baseline, shLDLR-treated sister cultures (to Figure 4D*i*) demonstrated a significant decrease in LDL-DyLight fluorescence compared to control (shLDLR +vehicle: 0.79-fold of control +vehicle ± 0.07). Control and scRNA-treated cultures exhibited expected significant increases in LDL-DyLight fluorescence after CT2074 or U18 treatment (Figure 4D*ii*, control in white bars, scRNA in gray bars). LDLR knockdown induced a significant decrease in CT2074-mediated LDL-DyLight fluorescence (Control +CT2074: 1.59-fold of vehicle ± 0.13; shLDLR +CT2074: 1.0-fold of vehicle ± 0.07). Similarly, U18-mediated LDL uptake was significantly decreased in shLDLR-treated cultures compared to control cultures treated with U18 (Control +U18: 3.11-fold of vehicle ± 0.28; shLDLR +U18: 1.66-fold of vehicle ± 0.11).

### 3.4 S2R modulator-induced LDL uptake requires TMEM97

To validate that the mechanism of action of the S2R modulators was indeed mediated through TMEM97, lentiviral shRNA vectors targeting TMEM97 (shTMEM97) were used to knockdown TMEM97 levels. In shTMEM97-treated cultures, TMEM97 mRNA was significantly decreased by 84% (0.16-fold of control ± 0.02) compared to control, while scrambled shRNA (scRNA) did not significantly change TMEM97 mRNA levels (Figure 5A). Similarly, shTMEM97 significantly decreased TMEM97 protein levels by 49% (mean ± 0.06) compared to control cultures (Figure 5B). We next asked whether this knockdown conferred a functional decrease in LDL uptake. At baseline, shTMEM97-treated cultures did not demonstrate significant changes in LDL uptake compared to control (Figure 5C). Control and scRNA-treated cultures exhibited expected significant increases in LDL-DyLight fluorescence after CT2074 or U18 treatment (Figure 5C, control in white bars, scRNA in gray bars). TMEM97 knockdown induced a significant decrease in LDL uptake mediated by CT2074 (Control +CT2074: 1.93-fold change of vehicle ± 0.14; shTMEM97 +CT2074: 1.46-fold of vehicle ± 0.09). Similarly, U18-mediated LDL uptake was significantly decreased in shTMEM97-treated cultures compared to control cultures treated with U18 (Control +U18: 3.14-fold change of vehicle ± 0.30; shTMEM97 +U18: 1.74-fold change of vehicle ± 0.08). The dependency of this effect on TMEM97 was further substantiated in an independent cell system: In wildtype HEK-293T cells, CT2074 (10 µM) induced a significant 1.47-fold (± 0.17) increase in LDL-DyLight levels compared to vehicle control, and effect that was obliterated in cells without TMEM97 (TMEM97-KO cells) (Supplemental Figure 3).

**Figure 5.**
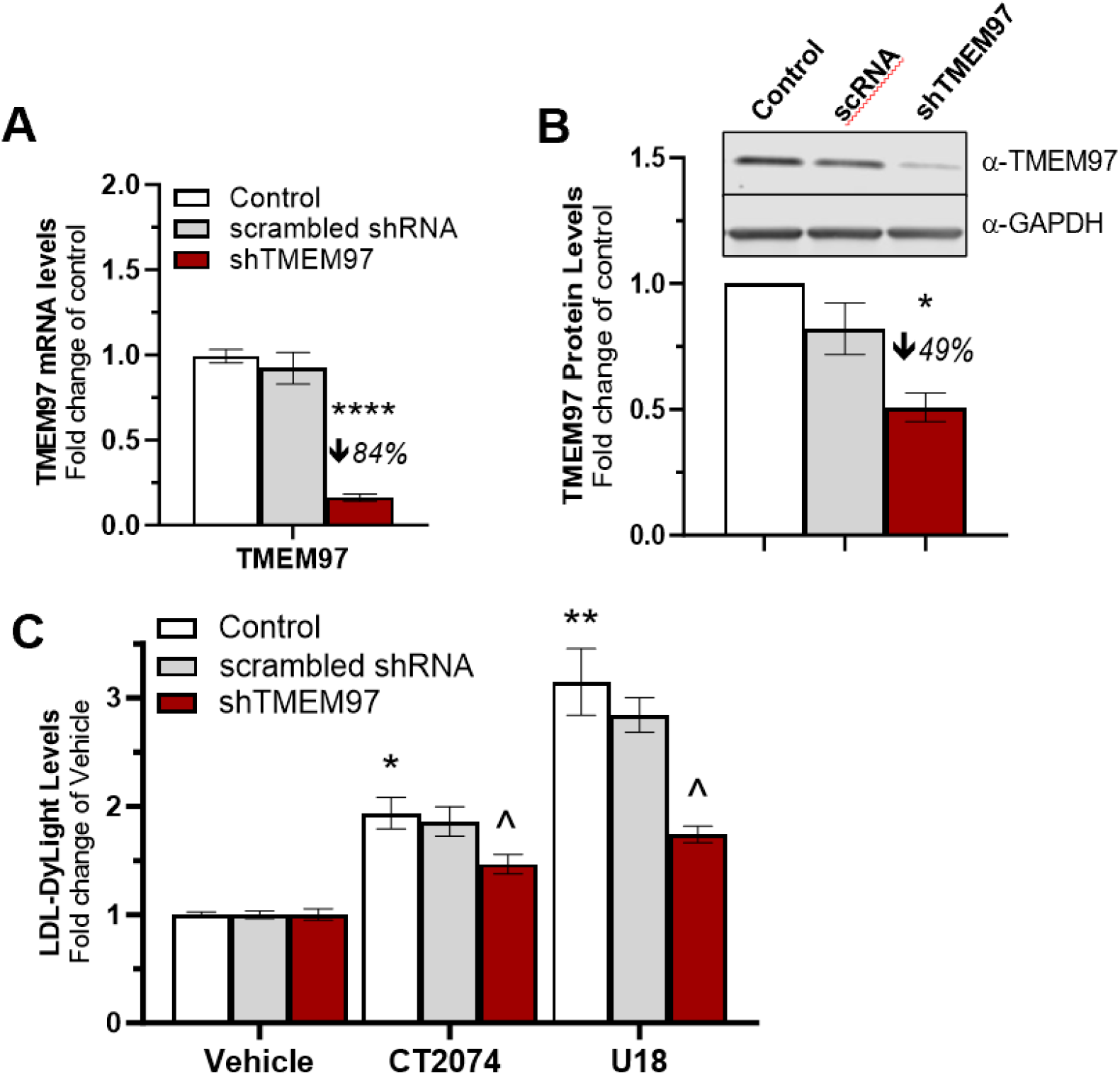
Increased LDL uptake via S2R modulators are dependent on TMEM97. TMEM97 levels in differentiated ARPE-19 cells treated with lentiviral vector carrying shRNA targeting TMEM97 (shTMEM97) or control vector (scRNA) were assessed by qRT-PCR **(A)** normalized to EIF4A2 and polybrene or by western blotting **(B)**, normalized to GAPDH and polybrene, mean ± SEM of N=3, percent TMEM97 knockdown indicated, significance assessed via one-way ANOVA, Dunnett’s multiple comparison test; p≤0.05 shTMEM97 vs control. **C)** Quantification of LDL-Dylight fluorescence after treatment with vehicle, CT2074 (10 µM) or U18 (5 µM) in control cells (white) or after treatment with scRNA (gray) or shTMEM97 (red) lentiviral vectors, normalized to vehicle, mean ± SEM of N=4, two-way ANOVA, Tukey’s multiple comparisons test: *p≤0.05, **p≤0.01 compound vs vehicle; ^p≤0.05, shTMEM97 vs control.

## 4 Discussion

The present study provides new evidence of a potential mechanism by which S2R modulators may impact retinal pigment epithelial cell function and characterizes this effect in a RPE cell-based model. In short, this study provides evidence that the S2R components and LDLR are expressed in differentiated ARPE-19 cells to add to the body of literature characterizing these proteins in mature RPE (Michelet et al., 2020; Ratnapriya et al., 2019; Shen et al., 2021a; Wang et al., 2023), and characterizes the mechanisms underlying basal LDL uptake as well as S2R- and U18-mediated LDL uptake in this model.

The S2R modulator CT1812 (zervimesine) was recently tested in a Ph2 clinical trial for geographic atrophy (GA) secondary to dry AMD (MAGNIFY, COG2201, NCT05893537). The interim analysis of participants in the MAGNIFY trial showed positive efficacy signals, wherein zervimesine-treated participants had 28.6% slower GA lesion growth on average compared to placebo (Cognition Therapeutics Inc., 2025). Preclinical studies further underscore the protective aspects of zervimesine and related proprietary S2R modulators. We have shown that treatment with a S2R modulator is protective in an in vivo glaucoma model (Donkor et al., 2024) and that S2R modulators applied to RPE cells in culture rescue POS trafficking deficits incurred by stressors relevant in dry AMD pathology (Lizama et al., 2025). The results described herein build upon the foundational understanding of S2R modulator mechanism of action that may be related to the protective effects of zervimesine observed in the MAGNIFY trial.

The ARPE-19 cell line has been used to facilitate understanding of dry AMD-relevant mechanisms, in that they are easy to maintain in culture for high-throughput studies, can be manipulated with genetic and pharmacological tools, and maintain the human genome to more accurately reflect human biology, unlike rodent or invertebrate models. However, undifferentiated ARPE-19 cultures do not exhibit the morphology nor key functions of native RPE cells, such as phagocytosis of POS. Appropriately differentiated ARPE-19 cells, as we and others have demonstrated (Hazim et al., 2019; Samuel et al., 2017), express the genes associated with visual cycle (*RPE65*), POS phagocytosis (*MERTK*), and terminal RPE maturation (*BEST1*) at high levels compared to undifferentiated ARPE-19 cells. We have also demonstrated herein that differentiated ARPE-19 cultures express the S2R components PGRMC1 and LDLR at higher levels than undifferentiated cells, which suggests that these receptors may play a key role in the functions of native human RPE. This is consistent with independent reports in ARPE-19 cells, in mature iPSC-derived RPE cells, and in primary RPE cells from mouse, and human postmortem RPE (Karlsson et al., 2021; Michelet et al., 2020; Ratnapriya et al., 2019; Shen et al., 2021a; Uhlén et al., 2015; Wang et al., 2023).

To interrogate S2R modulator mechanism of action, we developed an assay using differentiated ARPE-19 cells based on a key function of the S2R in regulating cellular levels of LDL-derived cholesterol (LDL-DyLight550 assay). Herein, we established that NPC1-specific cholesterol export inhibitor U18 (Lu et al., 2015) increases LDL uptake robustly in differentiated ARPE-19 cultures, which is strongly correlated with increased TMEM97 and LDLR transcript levels. These data corroborate a prior report showing that TMEM97 induction is dependent on sterol regulatory element-binding protein 2 (SREBP2) in the ARPE-19 cell line (Shen et al., 2021b). SREBP2 is the transcription factor coordinating cholesterol synthesis and uptake and controls LDLR levels (Horton et al., 2002). Using U18 as a positive control for LDL update in the RPE cultures, we then interrogated the effects of S2R modulators zervimesine, CT2074, and CT2168, effectively demonstrating that these modulators increase LDL uptake in RPE cells. We demonstrate that the LDL uptake by these S2R modulators is not due to off-target S1R binding, given that commercially-available S1R-specific small molecules do not induce LDL uptake up to 10 µM. These data support that LDL uptake is S2R – and not S1R – specific in RPE cells.

The S2R has been shown to be essential for intracellular homeostatic functions beyond LDLR-mediated lipid uptake. Other groups have shown that TMEM97 knockout RPE cells exhibit aberrant autophagy and lysosome dysfunction, as well as exacerbated oxidant-induced cell demise (Shen et al., 2021a). Autophagy, the regulated process of bulk protein and lipid sequestration and degradation, is critical to all cell types but is essential for RPE cells to perform the daily POS and lipoprotein degradation to support healthy retinas. Core autophagic machinery, namely LC3-associated vesicles and lysosomes, are required for the uptake and trafficking of POS (Keeling et al., 2020). We demonstrated that the same S2R modulators used herein that increase LDL uptake supported the healthy trafficking of POS in the presence of toxic Aβ protein or oxidative stress (Lizama et al., 2025). We also previously demonstrated that S2R modulators alter pathways related to vesicle trafficking and autophagy in transcriptomic analyses of RPE tissue from a mouse model of amyloid-beta accumulation (human APPsl transgenic mice) (Lizama et al., 2025). Interestingly, cholesterol biology and lipid metabolism were among the significantly altered pathways observed in zervimesine-treated animals (Lizama et al., 2025). Our results herein provided functional validation for the transcriptomic results by showing that zervimesine and other S2R modulators, alter LDL uptake. Due to the pivotal role LDLR plays in maintaining retinal and photoreceptor health (Chen et al., 2009; Sreekumar et al., 2022), it is tempting to speculate that the LDL trafficking induced by S2R modulators begets protection of POS trafficking. There is a wealth of literature to show that cholesterol, which is derived from LDL and from digested POS in RPE cells, is necessary to support the energetic requirements of cell functions, especially the fluidity of membranes comprising cellular vesicles and organelles (Lewandowski et al., 2022). Genetic knockdown/knockout studies have shown that LDLR deficiency results in photoreceptor degeneration, inflammation, and lipid accumulation (Chen et al., 2009; Sreekumar et al., 2022). Further investigation is warranted to dissect these potentially intersecting pathways of RPE lipid homeostasis, trafficking, and POS phagocytosis and processing.

We confirmed that S2R modulator mechanism of action in RPE cells is dependent on TMEM97 and LDLR using genetic knockdown and/or antibody-based tools. Using lentiviral vectors targeting TMEM97 or LDLR, basal protein levels were reduced by 49% and 32%, respectively. Interestingly, in vehicle-treated cultures, decreased levels of TMEM97 did not correspond to altered LDL uptake; on the other hand, decreased levels of LDLR (induced genetically or by antibody-mediated blockade of LDLR) significantly decreased LDL uptake in vehicle-treated cultures. These results suggest that under basal conditions, LDL uptake is chiefly LDLR-dependent and does not require TMEM97. However, a decrease of either TMEM97 or LDLR expression decreased S2R modulator- and U18-mediated LDL uptake. TMEM97 knockdown decreased the LDL uptake effect of CT2074, as well as the effect of zervimesine (preliminary data, not shown). These results validate S2R modulator engagement with TMEM97 to alter LDL uptake, as well as show that both TMEM97 and LDLR are necessary for this effect. We initially hypothesized that TMEM97 knockdown influences S2R modulator or U18 effects, more so than the control cultures, due to the 16h pretreatment that may induce changes in TMEM97 and/or LDLR expression levels, as concomitant or one-hour treatments did not alter LDL uptake (data not shown). Indeed, U18 treatment for 16h increased both TMEM97 and LDLR transcription. However, we did not find that S2R modulators induced corresponding increases in TMEM97 (data not shown) nor LDLR expression levels (Figure 4A), suggesting a distinct mechanism. In support, we observed that none of the S2R modulators at the maximal concentration used (10 µM) induced as much LDL uptake as U18 (< 2-fold change of control by S2R modulators, and >2-fold change of control by U18, 5 µM). This observation, combined with U18 being highly NPC1 specific (K_i_ = 0.03 µM) (Lu et al., 2015), suggests a potential NPC1-independent mechanism of S2R modulators for LDL uptake, although this remains to be investigated. Indeed, S2R interacts and shares a functional relationship with NPC1 (Bartz et al., 2009; Ebrahimi-Fakhari et al., 2015), indicating a pertinent downstream mechanism by which S2R can affect LDL trafficking. Elucidating the full scope of S2R’s involvement at different stages of LDL trafficking intracellularly warrants further research. To further underscore the point that TMEM97-mediated LDL uptake is context-dependent, we confirmed that HEK293T TMEM97 knockout cultures are unable to increase LDL uptake with S2R modulator treatment but, interestingly, are still able to increase LDL uptake in response to U18 treatment (wildtype: 2.51-fold ± 0.26 of vehicle; TMEM97-KO: 2.37-fold ± 0.23 of vehicle; Supplementary Figure 2). These data suggest that the effects of U18 may be cell-type specific, which is not surprising given that highly proliferative cells (such as HEK293T) have fluctuating demands for cholesterol and lipid composition as part of the cell cycle (Singh et al., 2013) and given cell type-specific regulation of LDLR, which has also been reported (Ye et al., 2014; Zheng et al., 2012).

We hypothesized that the mechanism for increased LDL uptake would be due to increased total protein levels of LDLR and/or TMEM97. Indeed, U18 increased LDL uptake and both LDLR and TMEM97 transcript levels in a concentration-dependent manner, in keeping with prior studies (Shen et al., 2021b). Surprisingly, while TMEM97 protein levels also increased in a U18 concentration-dependent manner up to 10 µM, LDLR protein levels did not show a significant increase at any concentration when assessed by western blot. We suspected that western blot was unable to capture the potential differences in subcellular pools of LDLR, which can be located in transport vesicles, endosomes, lysosomes, and at the cellular surface (Islam et al., 2022). Upon use of the LDLR-specific blocking antibody and fluorescence-based analyses, we observed increased levels of LDLR after U18 (as well as CT2074) treatment in non-permeabilized cells – that is, LDLR protein at the plasma membrane was increased after 16 hr treatment versus basal levels (Figure 4C*ii*), which may partially account for the observed elevated LDL uptake. Higher resolution imaging and differential centrifugation of cell lysate may allow further resolution of LDLR protein levels and discern differences in subcellular pools of LDLR after S2R modulator treatment. Additionally, assessing TMEM97 localization may be important to understanding S2R modulator mechanism of action in regulating lipid homeostasis, as we were unable to assess S2R plasma membrane levels like we could with LDLR given the lack of sophisticated antibody tools to target TMEM97 protein.

The abnormalities in complement activation seen in dry AMD have been the focus of much research and therapeutic development, and recent studies have identified other viable therapeutic targets. Dysfunctional lipid trafficking remains a major factor of dry AMD pathology, and while the field has dedicated attention to studying retinal lipid homeostasis and trafficking mechanisms, drug development efforts targeting these pathways have not yet materialized into approved clinical use for dry AMD. S2R modulators zervimesine and CT2074 have been shown to be blood-retina barrier penetrant and achieve adequate exposure levels in the retina with oral administration (Donkor et al., 2024; Lizama et al., 2025). The study herein represents the first demonstration of retina-penetrant small molecules targeting the S2R that can alter RPE LDL uptake. S2R modulators zervimesine, CT2074, and CT2168 induced increases in the LDL uptake assay, in keeping with our prior RPE POS trafficking studies (Lizama et al., 2025) showing that, while chemically-distinct, these S2R modulators exhibit similar pharmacology.

In sum, the results herein – taken together with the positive signal of efficacy of zervimesine in patients with dry AMD (MAGNIFY, COG2201, NCT05893537) – support development of S2R modulators to improve the dysfunctional lipid homeostasis observed in dry AMD. These results warrant further investigation of S2R’s role in lipid biology, given the data reported herein and published exploratory clinical trial biomarker data showing zervimesine modification of lipoprotein-related pathways in patients with AD ((B.N. Lizama et al., 2024; Britney N Lizama et al., 2024). While beyond the scope of this study, it would be interesting to observe effects of S2R modulators on lipid trafficking in an iPSC-derived RPE model, particularly those from donors diagnosed with dry AMD, to further resolve S2R modulator mechanism of action. Furthermore, due to the specialized functions of the RPE in regulating lipid and POS trafficking, it remains to be seen if U18 and S2R modulator mechanisms as reported herein are unique to RPE cells or broadly applicable to other cell types of the eye.

## Acknowledgments

The authors would like to thank Dr. Eunah Cho and Dr. Valentina Di Caro for feedback on the experimental design of the studies herein, as well as Ms. Mikayla Wilson and Ms. Nicole Knezovich for their technical support in cell culture and maintenance.

## Author Contributions

**BNL:**Conceptualization, Formal Analysis, Investigation, Methodology, Project Administration, Visualization, Writing – original draft, Writing – review and editing; **AR**: Formal analysis, Investigation, Methodology, Validation, Visualization, Writing – original draft, Writing – review and editing; **GL**: Resources, Writing – review and editing**; AOC:** Supervision, Writing – review and editing; **MEH**: Conceptualization, Visualization, Project Administration, Supervision; Writing – original draft, Writing – review and editing

## Supplementary Figure Legends

**Supplementary Figure 1:**
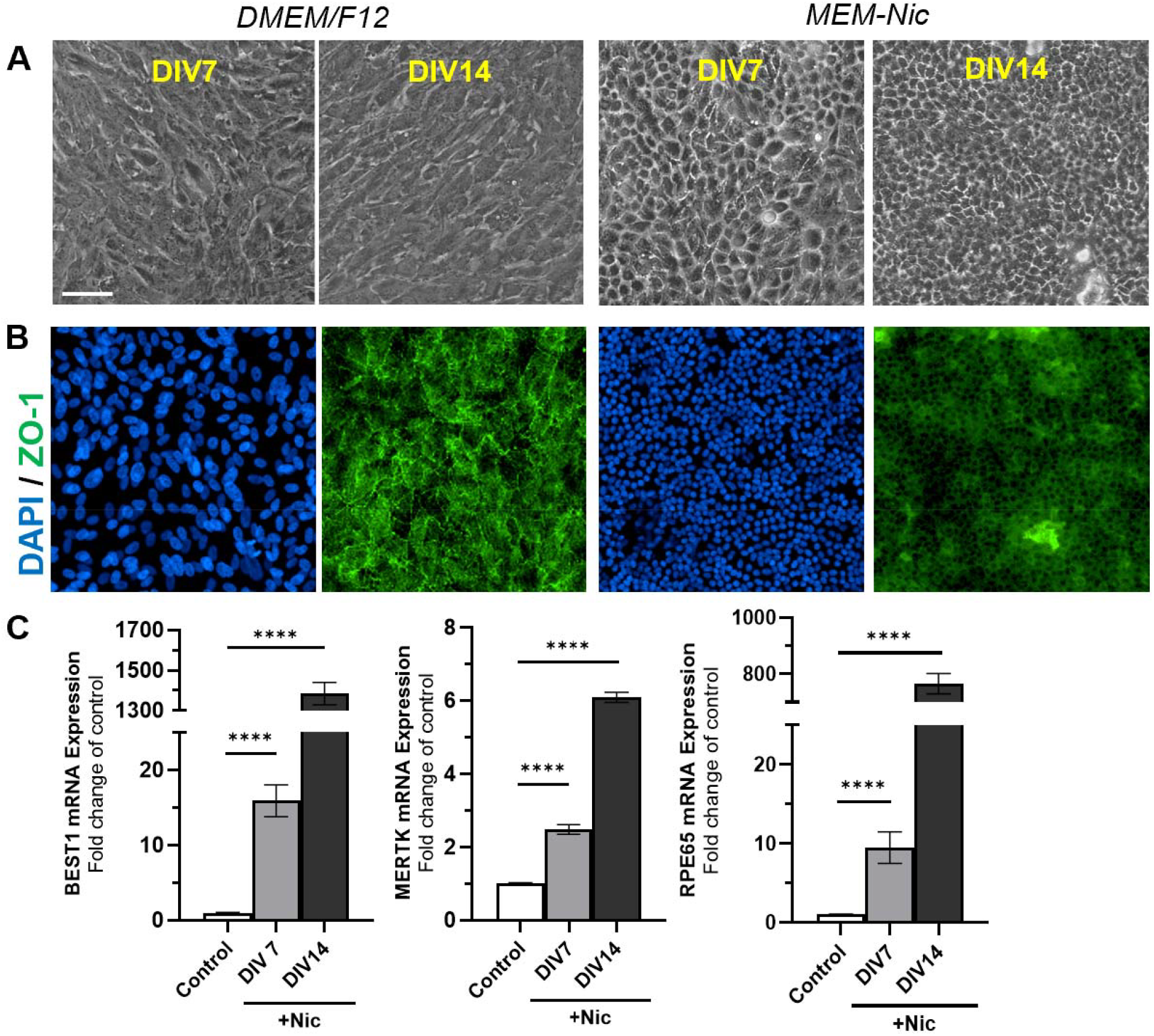
Differentiated ARPE-19 cells express markers of mature RPE. **A)** Brightfield images of ARPE-19 cells after 7 and 14 days of differentiation (DIV7 and DIV14 in MEM-Nic) compared to a control culture (DMEM:F12). 20x magnification, scale bar = 50 µm. **B)** Immunofluorescence staining showing the expression pattern of ZO-1 in undifferentiated ARPE19 cells (DMEM:F12) and ARPE19s differentiated with MEM-Nic (DIV14). 20x magnification, scale bar in A. **C)** qRT-PCR analysis of RPE-specific gene expression (*BEST1*, bestrophin-1; *MERTK*, Tyrosine-protein kinase Mer; *RPE65*, retinoid isomerohydrolase) after 7 and 14 days of differentiation in MEM-Nic, control conditions n=14, DIV7 conditions n=4, DIV14 conditions n=10, normalized to EIF4A2 + undifferentiated control, mean ± SEM, unpaired t-test. **** p≤0.0001 +Nic vs control.

**Supplementary Figure 2:**
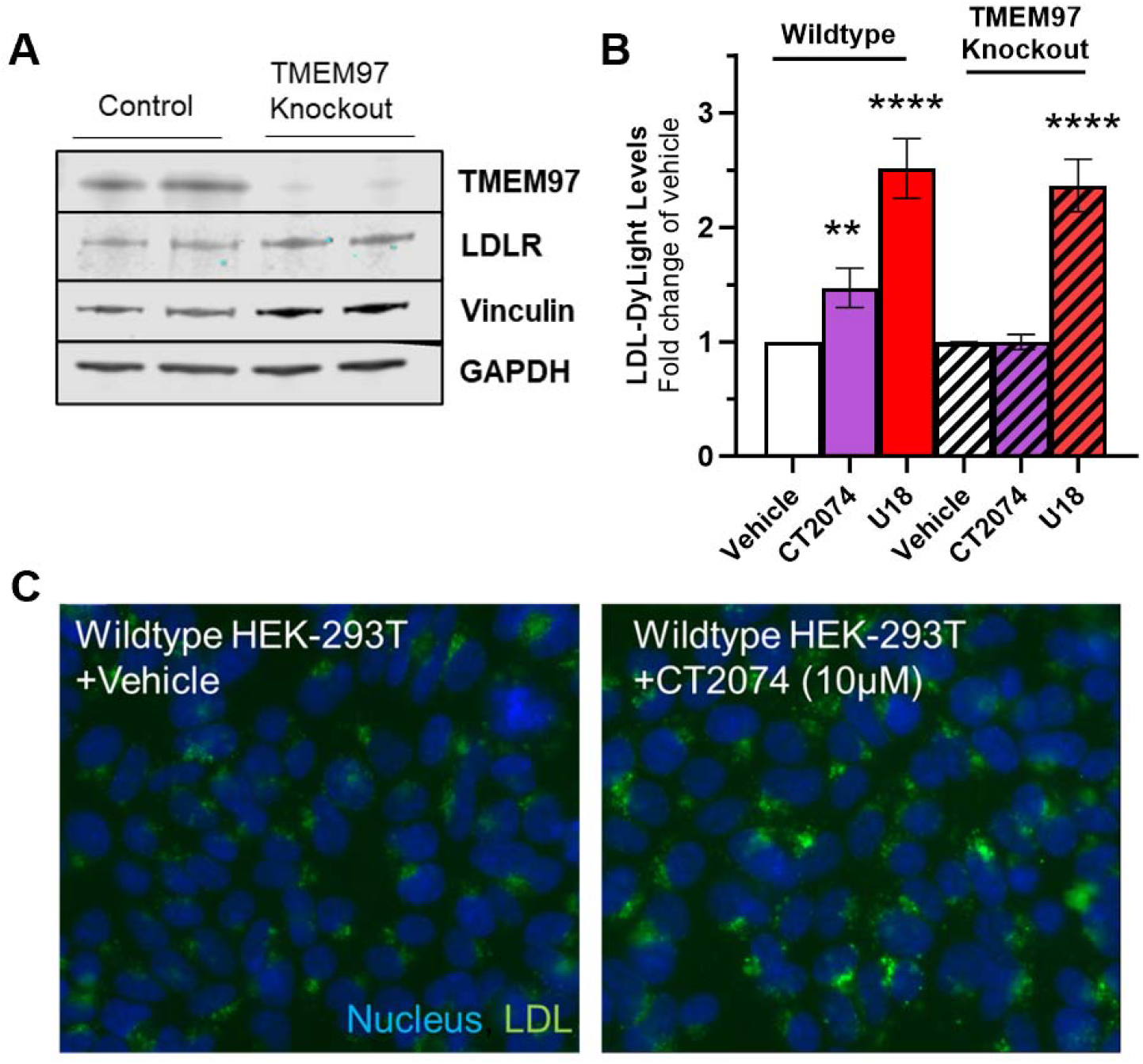
Increase in LDL uptake via S2R modulators is ablated in TMEM97 knockout cells. **A**) Western blots performed to confirm absence of TMEM97 in TMEM97-knockout HEK-293T cells and LDLR expression in wildtype and TMEM97 knockout cells. GAPDH and vinculin were assessed as loading controls. **B)** Quantification of LDL-Dylight assay fluorescence from wild-type or TMEM97-knockout HEK293 cells treated with CT2074 (10 μM) or U18666A (5 μM) (assessed by CX7 Spot Detector, Spot Total Intensity per Cell after treatment), N=6, normalized to vehicle, mean ± SEM, one-way ANOVA, Tukey’s multiple comparison test; **p≤0.01, ****p≤0.0001, treatment vs vehicle. **C)** Representative images of cells treated with vehicle or CT2074, labeled with LDL-DyLight (*green*) and Hoechst (*blue*).

## References

Arakawa, S., Takahashi, A., Ashikawa, K., Hosono, N., Aoi, T., Yasuda, M., Oshima, Y., Yoshida, S., Enaida, H., Tsuchihashi, T., Mori, K., Honda, S., Negi, A., Arakawa, A., Kadonosono, K., Kiyohara, Y., Kamatani, N., Nakamura, Y., Ishibashi, T., Kubo, M., 2011. Genome-wide association study identifies two susceptibility loci for exudative age-related macular degeneration in the Japanese population. Nat. Genet. 43, 1001–4. 10.1038/ng.938

Bartz, F., Kern, L., Erz, D., Zhu, M., Gilbert, D., Meinhof, T., Wirkner, U., Erfle, H., Muckenthaler, M., Pepperkok, R., Runz, H., 2009. Identification of Cholesterol-Regulating Genes by Targeted RNAi Screening. Cell Metab. 10, 63–75. 10.1016/j.cmet.2009.05.009

Chen, Y., Hu, Y., Moiseyev, G., Zhou, K.K., Chen, D., Ma, J., 2009. Photoreceptor degeneration and retinal inflammation induced by very low-density lipoprotein receptor deficiency. Microvasc. Res. 78, 119–127. 10.1016/j.mvr.2009.02.005

Cognition Therapeutics Inc., 2025. Cognition Therapeutics Reports Topline Results Showing Oral Zervimesine (CT1812) Reduced Lesion Growth in Phase 2 Study in Geographic Atrophy. Globe Newswire.

Curcio, C.A., 2018. Soft Drusen in Age-Related Macular Degeneration: Biology and Targeting Via the Oil Spill Strategies. Investig. Opthalmology Vis. Sci. 59, AMD160. 10.1167/iovs.18-24882

Curcio, C.A., Johnson, M., Rudolf, M., Huang, J.-D., 2011. The oil spill in ageing Bruch membrane. Br. J. Ophthalmol. 95, 1638–1645. 10.1136/bjophthalmol-2011-300344

Danzig, C.J., Khanani, A.M., Kaiser, P.K., Chang, M.A., Kovach, J.L., Lally, D.R., Rachitskaya, A., Sheth, V.S., Vajzovic, L., Clark, J., Tang, J., Zhu, L., Desai, D., Chakravarthy, U., 2024. Vision Loss Reduction with Avacincaptad Pegol for Geographic Atrophy. Ophthalmol. Retin. 8, 1052–1060. 10.1016/j.oret.2024.04.023

Donkor, N., Kiehlbauch, C.C., Pappenhagen, N., Look, G.C., Morgan, A.B., Shin, R., Hamby, M.E., Inman, D.M., 2024. Neuroprotective effect of Sigma-2 modulator CT2074 in a mouse model of ocular hypertension. Exp. Eye Res. 249, 110143. 10.1016/j.exer.2024.110143

Ebrahimi-Fakhari, D., Wahlster, L., Bartz, F., Werenbeck-Ueding, J., Praggastis, M., Zhang, J., Joggerst-Thomalla, B., Theiss, S., Grimm, D., Ory, D.S., Runz, H., 2015. Reduction of TMEM97 increases NPC1 protein levels and restores cholesterol trafficking in Niemann-pick type C1 disease cells. Hum. Mol. Genet. 25, 3588–3599. 10.1093/hmg/ddw204

Espinosa-Heidmann, D.G., Suner, I.J., Catanuto, P., Hernandez, E.P., Marin-Castano, M.E., Cousins, S.W., 2006. Cigarette Smoke–Related Oxidants and the Development of Sub-RPE Deposits in an Experimental Animal Model of Dry AMD. Investig. Opthalmology Vis. Sci. 47, 729. 10.1167/iovs.05-0719

Fleckenstein, M., Schmitz-Valckenberg, S., Chakravarthy, U., 2024. Age-Related Macular Degeneration. JAMA 331, 147. 10.1001/jama.2023.26074

Fritsche, L.G., Chen, W., Schu, M., Yaspan, B.L., Yu, Y., Thorleifsson, G., Zack, D.J., Arakawa, S., Cipriani, V., Ripke, S., Igo, R.P., Buitendijk, G.H.S., Sim, X., Weeks, D.E., Guymer, R.H., Merriam, J.E., Francis, P.J., Hannum, G., Agarwal, A., Armbrecht, A.M., Audo, I., Aung, T., Barile, G.R., Benchaboune, M., Bird, A.C., Bishop, P.N., Branham, K.E., Brooks, M., Brucker, A.J., Cade, W.H., Cain, M.S., Campochiaro, P.A., Chan, C.C., Cheng, C.Y., Chew, E.Y., Chin, K.A., Chowers, I., Clayton, D.G., Cojocaru, R., Conley, Y.P., Cornes, B.K., Daly, M.J., Dhillon, B., Edwards, A.O., Evangelou, E., Fagerness, J., Ferreyra, H.A., Friedman, J.S., Geirsdottir, A., George, R.J., Gieger, C., Gupta, N., Hagstrom, S.A., Harding, S.P., Haritoglou, C., Heckenlively, J.R., Holz, F.G., Hughes, G., Ioannidis, J.P.A., Ishibashi, T., Joseph, P., Jun, G., Kamatani, Y., Katsanis, N., N Keilhauer, C., Khan, J.C., Kim, I.K., Kiyohara, Y., Klein, B.E.K., Klein, R., Kovach, J.L., Kozak, I., Lee, C.J., Lee, K.E., Lichtner, P., Lotery, A.J., Meitinger, T., Mitchell, P., Mohand-Saïd, S., Moore, A.T., Morgan, D.J., Morrison, M.A., Myers, C.E., Naj, A.C., Nakamura, Y., Okada, Y., Orlin, A., Ortube, M.C., Othman, M.I., Pappas, C., Park, K.H., Pauer, G.J.T., Peachey, N.S., Poch, O., Priya, R.R., Reynolds, R., Richardson, A.J., Ripp, R., Rudolph, G., Ryu, E., Sahel, J.A., Schaumberg, D.A., Scholl, H.P.N., Schwartz, S.G., Scott, W.K., Shahid, H., Sigurdsson, H., Silvestri, G., Sivakumaran, T.A., Smith, R.T., Sobrin, L., Souied, E.H., Stambolian, D.E., Stefansson, H., Sturgill-Short, G.M., Takahashi, A., Tosakulwong, N., Truitt, B.J., Tsironi, E.E., Uitterlinden, A.G., Van Duijn, C.M., Vijaya, L., Vingerling, J.R., Vithana, E.N., Webster, A.R., Wichmann, H.E., Winkler, T.W., Wong, T.Y., Wright, A.F., Zelenika, D., Zhang, M., Zhao, L., Zhang, K., Klein, M.L., Hageman, G.S., Lathrop, G.M., Stefansson, K., Allikmets, R., Baird, P.N., Gorin, M.B., Wang, J.J., Klaver, C.C.W., Seddon, J.M., Pericak-Vance, M.A., Iyengar, S.K., Yates, J.R.W., Swaroop, A., Weber, B.H.F., Kubo, M., Deangelis, M.M., Léveillard, T., Thorsteinsdottir, U., Haines, J.L., Farrer, L.A., Heid, I.M., Abecasis, G.R., 2013. Seven new loci associated with age-related macular degeneration. Nat. Genet. 45, 433–439. 10.1038/ng.2578

Fritsche, L.G., Fariss, R.N., Stambolian, D., Abecasis, G.R., Curcio, C.A., Swaroop, A., 2014. Age-Related Macular Degeneration: Genetics and Biology Coming Together. Annu. Rev. Genomics Hum. Genet. 15, 151–171. 10.1146/annurev-genom-090413-025610

Fritsche, L.G., Igl, W., Bailey, J.N.C., Grassmann, F., Sengupta, S., Bragg-Gresham, J.L., Burdon, K.P., Hebbring, S.J., Wen, C., Gorski, M., Kim, I.K., Cho, D., Zack, D., Souied, E., Scholl, H.P.N., Bala, E., Lee, K.E., Hunter, D.J., Sardell, R.J., Mitchell, P., Merriam, J.E., Cipriani, V., Hoffman, J.D., Schick, T., Lechanteur, Y.T.E., Guymer, R.H., Johnson, M.P., Jiang, Y., Stanton, C.M., Buitendijk, G.H.S., Zhan, X., Kwong, A.M., Boleda, A., Brooks, M., Gieser, L., Ratnapriya, R., Branham, K.E., Foerster, J.R., Heckenlively, J.R., Othman, M.I., Vote, B.J., Liang, H.H., Souzeau, E., McAllister, I.L., Isaacs, T., Hall, J., Lake, S., Mackey, D.A., Constable, I.J., Craig, J.E., Kitchner, T.E., Yang, Z., Su, Z., Luo, H., Chen, D., Ouyang, H., Flagg, K., Lin, D., Mao, G., Ferreyra, H., Stark, K., von Strachwitz, C.N., Wolf, A., Brandl, C., Rudolph, G., Olden, M., Morrison, M.A., Morgan, D.J., Schu, M., Ahn, J., Silvestri, G., Tsironi, E.E., Park, K.H., Farrer, L.A., Orlin, A., Brucker, A., Li, M., Curcio, C.A., Mohand-Saïd, S., Sahel, J.-A., Audo, I., Benchaboune, M., Cree, A.J., Rennie, C.A., Goverdhan, S. V, Grunin, M., Hagbi-Levi, S., Campochiaro, P., Katsanis, N., Holz, F.G., Blond, F., Blanché, H., Deleuze, J.-F., Igo, R.P., Truitt, B., Peachey, N.S., Meuer, S.M., Myers, C.E., Moore, E.L., Klein, R., Hauser, M.A., Postel, E.A., Courtenay, M.D., Schwartz, S.G., Kovach, J.L., Scott, W.K., Liew, G., Tan, A.G., Gopinath, B., Merriam, J.C., Smith, R.T., Khan, J.C., Shahid, H., Moore, A.T., McGrath, J.A., Laux, R., Brantley, M.A., Agarwal, A., Ersoy, L., Caramoy, A., Langmann, T., Saksens, N.T.M., de Jong, E.K., Hoyng, C.B., Cain, M.S., Richardson, A.J., Martin, T.M., Blangero, J., Weeks, D.E., Dhillon, B., van Duijn, C.M., Doheny, K.F., Romm, J., Klaver, C.C.W., Hayward, C., Gorin, M.B., Klein, M.L., Baird, P.N., den Hollander, A.I., Fauser, S., Yates, J.R.W., Allikmets, R., Wang, J.J., Schaumberg, D.A., Klein, B.E.K., Hagstrom, S.A., Chowers, I., Lotery, A.J., Léveillard, T., Zhang, K., Brilliant, M.H., Hewitt, A.W., Swaroop, A., Chew, E.Y., Pericak-Vance, M.A., DeAngelis, M., Stambolian, D., Haines, J.L., Iyengar, S.K., Weber, B.H.F., Abecasis, G.R., Heid, I.M., 2016. A large genome-wide association study of age-related macular degeneration highlights contributions of rare and common variants. Nat. Genet. 48, 134–43. 10.1038/ng.3448

Go, G.-W., Mani, A., 2012. Low-density lipoprotein receptor (LDLR) family orchestrates cholesterol homeostasis. Yale J. Biol. Med. 85, 19–28.

Gordiyenko, N., Campos, M., Lee, J.W., Fariss, R.N., Sztein, J., Rodriguez, I.R., 2004. RPE Cells Internalize Low-Density Lipoprotein (LDL) and Oxidized LDL (oxLDL) in Large Quantities In Vitro and In Vivo. Investig. Opthalmology Vis. Sci. 45, 2822. 10.1167/iovs.04-0074

Hazim, R.A., Volland, S., Yen, A., Burgess, B.L., Williams, D.S., 2019. Rapid differentiation of the human RPE cell line, ARPE-19, induced by nicotinamide. Exp. Eye Res. 179, 18–24. 10.1016/j.exer.2018.10.009

Helgason, H., Sulem, P., Duvvari, M.R., Luo, H., Thorleifsson, G., Stefansson, H., Jonsdottir, I., Masson, G., Gudbjartsson, D.F., Walters, G.B., Magnusson, O.T., Kong, A., Rafnar, T., Kiemeney, L.A., Schoenmaker-Koller, F.E., Zhao, L., Boon, C.J.F., Song, Y., Fauser, S., Pei, M., Ristau, T., Patel, S., Liakopoulos, S., van de Ven, J.P.H., Hoyng, C.B., Ferreyra, H., Duan, Y., Bernstein, P.S., Geirsdottir, A., Helgadottir, G., Stefansson, E., den Hollander, A.I., Zhang, K., Jonasson, F., Sigurdsson, H., Thorsteinsdottir, U., Stefansson, K., 2013. A rare nonsynonymous sequence variant in C3 is associated with high risk of age-related macular degeneration. Nat. Genet. 45, 1371–1374. 10.1038/ng.2740

Horton, J.D., Goldstein, J.L., Brown, M.S., 2002. SREBPs: activators of the complete program of cholesterol and fatty acid synthesis in the liver. J. Clin. Invest. 109, 1125–1131. 10.1172/JCI200215593

Huisingh, C., McGwin, G., Neely, D., Zarubina, A., Clark, M., Zhang, Y., Curcio, C.A., Owsley, C., 2016. The Association Between Subretinal Drusenoid Deposits in Older Adults in Normal Macular Health and Incident Age-Related Macular Degeneration. Investig. Opthalmology Vis. Sci. 57, 739. 10.1167/iovs.15-18316

Islam, M.M., Hlushchenko, I., Pfisterer, S.G., 2022. Low-Density Lipoprotein Internalization, Degradation and Receptor Recycling Along Membrane Contact Sites. Front. Cell Dev. Biol. 10. 10.3389/fcell.2022.826379

Izzo, N.J., Staniszewski, A., To, L., Fa, M., Teich, A.F., Saeed, F., Wostein, H., Walko, T., Vaswani, A., Wardius, M., Syed, Z., Ravenscroft, J., Mozzoni, K., Silky, C., Rehak, C., Yurko, R., Finn, P., Look, G., Rishton, G., Safferstein, H., Miller, M., Johanson, C., Stopa, E., Windisch, M., Hutter-Paier, B., Shamloo, M., Arancio, O., LeVine, H., Catalano, S.M., 2014a. Alzheimer’s therapeutics targeting amyloid beta 1-42 oligomers I: Abeta 42 oligomer binding to specific neuronal receptors is displaced by drug candidates that improve cognitive deficits. PLoS One 9. 10.1371/journal.pone.0111898

Izzo, N.J., Xu, J., Zeng, C., Kirk, M.J., Mozzoni, K., Silky, C., Rehak, C., Yurko, R., Look, G., Rishton, G., Safferstein, H., Cruchaga, C., Goate, A., Cahill, M.A., Arancio, O., Mach, R.H., Craven, R., Head, E., LeVine, H., Spires-Jones, T.L., Catalano, S.M., 2014b. Alzheimer’s Therapeutics Targeting Amyloid Beta 1–42 Oligomers II: Sigma-2/PGRMC1 Receptors Mediate Abeta 42 Oligomer Binding and Synaptotoxicity. PLoS One 9, e111899. 10.1371/journal.pone.0111899

Izzo, N.J., Yuede, C.M., LaBarbera, K.M., Limegrover, C.S., Rehak, C., Yurko, R., Waybright, L., Look, G., Rishton, G., Safferstein, H., Hamby, M.E., Williams, C., Sadlek, K., Edwards, H.M., Davis, C.S., Grundman, M., Schneider, L.S., DeKosky, S.T., Chelsky, D., Pike, I., Henstridge, C., Blennow, K., Zetterberg, H., LeVine, H., Spires-Jones, T.L., Cirrito, J.R., Catalano, S.M., 2021. Preclinical and clinical biomarker studies of CT1812: A novel approach to Alzheimer’s disease modification. Alzheimer’s Dement. 17, 1365–1382. 10.1002/alz.12302

Jaffe, G.J., Westby, K., Csaky, K.G., Monés, J., Pearlman, J.A., Patel, S.S., Joondeph, B.C., Randolph, J., Masonson, H., Rezaei, K.A., 2021. C5 Inhibitor Avacincaptad Pegol for Geographic Atrophy Due to Age-Related Macular Degeneration. Ophthalmology 128, 576–586. 10.1016/j.ophtha.2020.08.027

Kailani, Z., Mihalache, A., Popovic, M.M., Kertes, P.J., Muni, R.H., 2025. Ocular Adverse Events Associated with Pegcetacoplan and Avacincaptad Pegol for Geographic Atrophy: A Population-Based Pharmacovigilance Study. Am. J. Ophthalmol. 276, 252–260. 10.1016/j.ajo.2025.04.022

Karlsson, M., Zhang, C., Méar, L., Zhong, W., Digre, A., Katona, B., Sjöstedt, E., Butler, L., Odeberg, J., Dusart, P., Edfors, F., Oksvold, P., von Feilitzen, K., Zwahlen, M., Arif, M., Altay, O., Li, X., Ozcan, M., Mardonoglu, A., Fagerberg, L., Mulder, J., Luo, Y., Ponten, F., Uhlén, M., Lindskog, C., 2021. A single–cell type transcriptomics map of human tissues. Sci. Adv. 7, 1–9. 10.1126/sciadv.abh2169

Keeling, E., Chatelet, D.S., Johnston, D.A., Page, A., Tumbarello, D.A., Lotery, A.J., Ratnayaka, J.A., 2019. Oxidative Stress and Dysfunctional Intracellular Traffic Linked to an Unhealthy Diet Results in Impaired Cargo Transport in the Retinal Pigment Epithelium (RPE). Mol. Nutr. Food Res. 63, 1–18. 10.1002/mnfr.201800951

Keeling, E., Culling, A.J., Johnston, D.A., Chatelet, D.S., Page, A., Tumbarello, D.A., Lotery, A.J., Ratnayaka, J.A., 2020. An In-Vitro Cell Model of Intracellular Protein Aggregation Provides Insights into RPE Stress Associated with Retinopathy. Int. J. Mol. Sci. 21, 6647. 10.3390/ijms21186647

Keeling, E., Lotery, A., Tumbarello, D., Ratnayaka, J., 2018. Impaired Cargo Clearance in the Retinal Pigment Epithelium (RPE) Underlies Irreversible Blinding Diseases. Cells 7, 16. 10.3390/cells7020016

Liao, D.S., Grossi, F. V., El Mehdi, D., Gerber, M.R., Brown, D.M., Heier, J.S., Wykoff, C.C., Singerman, L.J., Abraham, P., Grassmann, F., Nuernberg, P., Weber, B.H.F., Deschatelets, P., Kim, R.Y., Chung, C.Y., Ribeiro, R.M., Hamdani, M., Rosenfeld, P.J., Boyer, D.S., Slakter, J.S., Francois, C.G., 2020. Complement C3 Inhibitor Pegcetacoplan for Geographic Atrophy Secondary to Age-Related Macular Degeneration: A Randomized Phase 2 Trial. Ophthalmology 127, 186–195. 10.1016/j.ophtha.2019.07.011

Liao, D.S., Metlapally, R., Joshi, P., 2022. Pegcetacoplan Treatment for Geographic Atrophy Due to Age-Related Macular Degeneration: A Plain Language Summary of the FILLY Study. Immunotherapy 14, 995–1006. 10.2217/imt-2022-0078

Lizama, B.N., Kahle, J., Catalano, S.M., Caggiano, A.O., Grundman, M., Hamby, M.E., 2023. Sigma-2 Receptors— From Basic Biology to Therapeutic Target: A Focus on Age-Related Degenerative Diseases. Int. J. Mol. Sci. 10.3390/ijms24076251

Lizama, B.N., Keeling, E., Cho, E., Malagise, E.M., Knezovich, N., Waybright, L., Watto, E., Look, G., Di Caro, V., Caggiano, A.O., Ratnayaka, J.A., Hamby, M.E., 2025. Sigma-2 receptor modulator CT1812 alters key pathways and rescues retinal pigment epithelium (RPE) functional deficits associated with dry age-related macular degeneration (AMD). Sci. Rep. 15, 4256. 10.1038/s41598-025-87921-9

Lizama, B.N., North, H.A., Pandey, K., Williams, C., Duong, D., Cho, E., Di Caro, V., Ping, L., Blennow, K., Zetterberg, H., Lah, J., Levey, A.I., Grundman, M., Caggiano, A.O., Seyfried, N.T., Hamby, M.E., 2024. An interim exploratory proteomics biomarker analysis of a phase 2 clinical trial to assess the impact of CT1812 in Alzheimer’s disease. Neurobiol. Dis. 199, 106575. 10.1016/j.nbd.2024.106575

Lizama, Britney N, Williams, C., North, H.A., Pandey, K., Duong, D., Di Caro, V., Mecca, A.P., Blennow, K., Zetterberg, H., Levey, A.I., Grundman, M., van Dyck, C.H., Caggiano, A.O., Seyfried, N.T., Hamby, M.E., 2024. CT1812 biomarker signature from a meta-analysis of CSF proteomic findings from two Phase 2 clinical trials in Alzheimer’s disease. Alzheimers. Dement. 20, 6860–6880. 10.1002/alz.14152

Lu, F., Liang, Q., Abi-Mosleh, L., Das, A., De Brabander, J.K., Goldstein, J.L., Brown, M.S., 2015. Identification of NPC1 as the target of U18666A, an inhibitor of lysosomal cholesterol export and Ebola infection. Elife 4. 10.7554/eLife.12177

Lynn, S.A., Pandi, S.P.S., Sanchez-Bretano, A., Muir, A.-M., Parker, L., Chatelet, D.S., Newall, T., Scott, J.A., Keeling, E., Smyth, N.R., Self, J.E., Lotery, A.J., Lee, H., Ratnayaka, J.A., 2025. A longitudinal study of the 5xFAD mouse retina delineates Amyloid beta (Aβ)-mediated retinal pathology from age-related changes. Alzheimers. Res. Ther. 17, 136. 10.1186/s13195-025-01784-w

Michelet, F., Balasankar, A., Teo, N., Stanton, L.W., Singhal, S., 2020. Rapid generation of purified human RPE from pluripotent stem cells using 2D cultures and lipoprotein uptake-based sorting. Stem Cell Res. Ther. 11, 47. 10.1186/s13287-020-1568-3

Nadeem, A., Malik, I.A., Shariq, F., Afridi, E.K., Taha, M., Raufi, N., Naveed, A.K., Iqbal, J., Habte, A., 2023. Advancements in the treatment of geographic atrophy: focus on pegcetacoplan in age-related macular degeneration. Ann. Med. Surg. 85, 6067–6077. 10.1097/MS9.0000000000001466

Neale, B.M., Fagerness, J., Reynolds, R., Sobrin, L., Parker, M., Raychaudhuri, S., Tan, P.L., Oh, E.C., Merriam, J.E., Souied, E., Bernstein, P.S., Li, B., Frederick, J.M., Zhang, K., Brantley, M.A., Lee, A.Y., Zack, D.J., Campochiaro, B., Campochiaro, P., Ripke, S., Smith, R.T., Barile, G.R., Katsanis, N., Allikmets, R., Daly, M.J., Seddon, J.M., 2010. Genome-wide association study of advanced age-related macular degeneration identifies a role of the hepatic lipase gene (LIPC). Proc. Natl. Acad. Sci. 107, 7395–7400. 10.1073/pnas.0912019107

Pikuleva, I.A., Curcio, C.A., 2014. Cholesterol in the retina: The best is yet to come. Prog. Retin. Eye Res. 41, 64– 89. 10.1016/j.preteyeres.2014.03.002

Ratnapriya, R., Sosina, O.A., Starostik, M.R., Kwicklis, M., Kapphahn, R.J., Fritsche, L.G., Walton, A., Arvanitis, M., Gieser, L., Pietraszkiewicz, A., Montezuma, S.R., Chew, E.Y., Battle, A., Abecasis, G.R., Ferrington, D.A., Chatterjee, N., 2019. Retinal transcriptome and eQTL analyses identify genes associated with age-related macular degeneration. Nat Genet 51, 606–610. 10.1038/s41588-019-0351-9.Retinal

Raychaudhuri, S., Iartchouk, O., Chin, K., Tan, P.L., Tai, A.K., Ripke, S., Gowrisankar, S., Vemuri, S., Montgomery, K., Yu, Y., Reynolds, R., Zack, D.J., Campochiaro, B., Campochiaro, P., Katsanis, N., Daly, M.J., Seddon, J.M., 2011. A rare penetrant mutation in CFH confers high risk of age-related macular degeneration. Nat. Genet. 43, 1232–1236. 10.1038/ng.976

Riad, A., Lengyel-Zhand, Z., Zeng, C., Weng, C.-C., Lee, V.M.-Y., Trojanowski, J.Q., Mach, R.H., 2020. The Sigma-2 Receptor/TMEM97, PGRMC1, and LDL Receptor Complex Are Responsible for the Cellular Uptake of Aβ42 and Its Protein Aggregates. Mol. Neurobiol. 57, 3803–3813. 10.1007/s12035-020-01988-1

Riad, A., Zeng, C., Weng, C.-C., Winters, H., Xu, K., Makvandi, M., Metz, T., Carlin, S., Mach, R.H., 2018. Sigma-2 Receptor/TMEM97 and PGRMC-1 Increase the Rate of Internalization of LDL by LDL Receptor through the Formation of a Ternary Complex. Sci. Rep. 8, 16845. 10.1038/s41598-018-35430-3

Rudolf, M., Ivandic, B., Winkler, J., Schmidt-Erfurth, U., 2004. Accumulation of lipid particles in the rupture membrane of LDL receptor-deficient mice as a model for age-related macular degeneration. Der Ophthalmol. 101, 715–719. 10.1007/s00347-003-0942-8

Samuel, W., Jaworski, C., Postnikova, O.A., Kutty, R.K., Duncan, T., Tan, L.X., Poliakov, E., Lakkaraju, A., Redmond, T.M., 2017. Appropriately differentiated ARPE-19 cells regain phenotype and gene expression profiles similar to those of native RPE cells. Mol. Vis. 23, 60–89.

Seddon, J.M., Yu, Y., Miller, E.C., Reynolds, R., Tan, P.L., Gowrisankar, S., Goldstein, J.I., Triebwasser, M., Anderson, H.E., Zerbib, J., Kavanagh, D., Souied, E., Katsanis, N., Daly, M.J., Atkinson, J.P., Raychaudhuri, S., 2013. Rare variants in CFI, C3 and C9 are associated with high risk of advanced age-related macular degeneration. Nat. Genet. 45, 1366–1370. 10.1038/ng.2741

Shen, H., Li, J., Heisler-Taylor, T., Makin, R., Yang, H., Mavlyutov, T.A., Gelfand, B., Cebulla, C.M., Guo, L.-W., 2021a. TMEM97 ablation aggravates oxidant-induced retinal degeneration. Cell. Signal. 86, 110078. 10.1016/j.cellsig.2021.110078

Shen, H., Li, J., Xie, X., Yang, H., Zhang, M., Wang, B., Kent, K.C., Plutzky, J., Guo, L.-W., 2021b. BRD2 regulation of sigma-2 receptor upon cholesterol deprivation. Life Sci. Alliance 4, e201900540. 10.26508/lsa.201900540

Singh, P., Saxena, R., Srinivas, G., Pande, G., Chattopadhyay, A., 2013. Cholesterol Biosynthesis and Homeostasis in Regulation of the Cell Cycle. PLoS One 8, e58833. 10.1371/journal.pone.0058833

Śpiewak, D., Drzyzga, Ł., Dorecka, M., Wyględowska-Promieńska, D., 2024. Summary of the Therapeutic Options for Patients with Dry and Neovascular AMD. J. Clin. Med. 13, 4227. 10.3390/jcm13144227

Sreekumar, P.G., Su, F., Spee, C., Araujo, E., Nusinowitz, S., Reddy, S.T., Kannan, R., 2022. Oxidative Stress and Lipid Accumulation Augments Cell Death in LDLR-Deficient RPE Cells and Ldlr−/− Mice. Cells 12, 43. 10.3390/cells12010043

Tserentsoodol, N., Sztein, J., Campos, M., Gordiyenko, N. V, Fariss, R.N., Lee, J.W., Fliesler, S.J., Rodriguez, I.R., 2006. Uptake of cholesterol by the retina occurs primarily via a low density lipoprotein receptor-mediated process. Mol. Vis. 12, 1306–18.

Uhlén, M., Fagerberg, L., Hallström, B.M., Lindskog, C., Oksvold, P., Mardinoglu, A., Sivertsson, Å., Kampf, C., Sjöstedt, E., Asplund, A., Olsson, I., Edlund, K., Lundberg, E., Navani, S., Szigyarto, C.A.-K., Odeberg, J., Djureinovic, D., Takanen, J.O., Hober, S., Alm, T., Edqvist, P.-H., Berling, H., Tegel, H., Mulder, J., Rockberg, J., Nilsson, P., Schwenk, J.M., Hamsten, M., von Feilitzen, K., Forsberg, M., Persson, L., Johansson, F., Zwahlen, M., von Heijne, G., Nielsen, J., Pontén, F., 2015. Tissue-based map of the human proteome. Science (80-.). 347. 10.1126/science.1260419

van Dyck, C.H., Mecca, A.P., O’Dell, R.S., Bartlett, H.H., Diepenbrock, N.G., Huang, Y., Hamby, M.E., Grundman, M., Catalano, S.M., Caggiano, A.O., Carson, R.E., 2024. A pilot study to evaluate the effect of CT1812 treatment on synaptic density and other biomarkers in Alzheimer’s disease. Alzheimers. Res. Ther. 16, 20. 10.1186/s13195-024-01382-2

Veerappan, M., El-Hage-Sleiman, A.-K.M., Tai, V., Chiu, S.J., Winter, K.P., Stinnett, S.S., Hwang, T.S., Hubbard, G.B., Michelson, M., Gunther, R., Wong, W.T., Chew, E.Y., Toth, C.A., Toth, C.A., Wong, W., Hwang, T., Hubbard, G.B., Srivastava, S., McCall, M., Winter, K., Sarin, N., Hall, K., McCollum, P., Curtis, L., Schuman, S., Chiu, S.J., Farsiu, S., Tai, V., Sevilla, M., Harrington, C., Gunther, R., Tran-Viet, D., Folgar, F., Yuan, E., Clemons, T., Harrington, M., Chew, E., 2016. Optical Coherence Tomography Reflective Drusen Substructures Predict Progression to Geographic Atrophy in Age-related Macular Degeneration. Ophthalmology 123, 2554–2570. 10.1016/j.ophtha.2016.08.047

Vijverberg, E., de Haan, W., Scheijbeler, E., Hamby, M.E., Catalano, S., Scheltens, P., Grundman, M., Caggiano, A.O., 2024. A Pilot Electroencephalography Study of the Effect of CT1812 Treatment on Synaptic Activity in Patients with Mild to Moderate Alzheimer’s Disease. J. Prev. Alzheimer’s Dis. 11, 1809–1817. 10.14283/jpad.2024.154

Vijverberg, E.G.B., Scharre, D.W., Woodward, M., Catalano, S., Hamby, M.E., Grundman, M., Morgan, R.E., Iaci, J., Devins, T., Caggiano, A.O., 2025. Randomized, Double-Blind Study of Zervimesine in Mild to Moderate Alzheimer’s Disease. Submitted

Wang, J.-H., Urrutia-Cabrera, D., Lees, J.G., Mesa Mora, S., Nguyen, T., Hung, S.S.C., Hewitt, A.W., Lim, S.Y., Edwards, T.L., Wong, R.C.B., 2023. Development of a CRISPRi Human Retinal Pigmented Epithelium Model for Functional Study of Age-Related Macular Degeneration Genes. Int. J. Mol. Sci. 24, 3417. 10.3390/ijms24043417

Yamada, Y., Tian, J., Yang, Y., Cutler, R.G., Wu, T., Telljohann, R.S., Mattson, M.P., Handa, J.T., 2008. Oxidized low density lipoproteins induce a pathologic response by retinal pigmented epithelial cells. J. Neurochem. 105, 1187–1197. 10.1111/j.1471-4159.2008.05211.x

Yan, Q., Ding, Y., Liu, Yi, Sun, T., Fritsche, L.G., Clemons, T., Ratnapriya, R., Klein, M.L., Cook, R.J., Liu, Yu, Fan, R., Wei, L., Abecasis, G.R., Swaroop, A., Chew, E.Y., Weeks, D.E., Chen, W., 2018. Genome-wide analysis of disease progression in age-related macular degeneration. Hum. Mol. Genet. 27, 929–940. 10.1093/hmg/ddy002

Ye, Q., Lei, H., Fan, Z., Zheng, W., Zheng, S., 2014. Difference in LDL Receptor Feedback Regulation in Macrophages and Vascular Smooth Muscle Cells: Foam Cell Transformation Under Inflammatory Stress. Inflammation 37, 555–565. 10.1007/s10753-013-9769-x

Zheng, W., Reem, R.E., Omarova, S., Huang, S., DiPatre, P.L., Charvet, C.D., Curcio, C.A., Pikuleva, I.A., 2012. Spatial Distribution of the Pathways of Cholesterol Homeostasis in Human Retina. PLoS One 7, e37926. 10.1371/journal.pone.0037926

